# Proteomic analysis identifies the E3 ubiquitin ligase Pdzrn3 as a regulatory target of Wnt5a-Ror signaling

**DOI:** 10.1101/2020.06.22.163790

**Authors:** Sara E. Konopelski Snavely, Michael W. Susman, Ryan C. Kunz, Jia Tan, Srisathya Srinivasan, Michael D. Cohen, Kyoko Okada, Helen Lamb, Shannon S. Choi, Edith P. Karuna, Michael K. Scales, Steven P. Gygi, Michael E. Greenberg, Hsin-Yi Henry Ho

## Abstract

Wnt5a-Ror signaling is a conserved pathway that regulates morphogenetic processes during vertebrate development, but its downstream signaling events remain poorly understood. Through a large-scale proteomic screen in mouse embryonic fibroblasts, we identified the E3 ubiquitin ligase Pdzrn3 as a regulatory target of the Wnt5a-Ror pathway. Upon pathway activation, Pdzrn3 is degraded in a β-catenin-independent, ubiquitin-proteasome system-dependent manner. We developed a flow cytometry-based reporter to monitor Pdzrn3 abundance and delineated a signaling cascade involving Frizzled, Dishevelled, CK1, and GSK3 that regulates Pdzrn3 stability. Epistatically, Pdzrn3 is regulated independently of Kif26b, another Wnt5a-Ror effector. Wnt5a-dependent degradation of Pdzrn3 requires phosphorylation of three conserved amino acids within its C-terminal LNX3H domain, which acts as a bona fide Wnt5a-responsive element. Importantly, this phospho-dependent degradation is essential for Wnt5a-Ror modulation of cell migration. Collectively, this work establishes a new Wnt5a-Ror cell morphogenetic cascade involving Pdzrn3 phosphorylation and degradation.

## Introduction

Embryonic development in vertebrates is a highly stereotyped and coordinated process that depends on a handful of core signaling pathways. One major mode of signaling involves Wnt ligands, a diverse and highly conserved family of glycoproteins that signal in many spatiotemporal contexts, including tissue specification and tissue morphogenesis in addition to tissue homeostasis in adult organisms (Clevers & Nusse, 2012; Nusse & Varmus, 2012; Steinhart & Angers, 2018). Thus, Wnts play unique and critical roles in both developing and adult organisms.

Traditionally, Wnt pathways have been classified as either canonical or non-canonical. Canonical Wnt signaling utilizes β-catenin as a transcriptional co-activator to regulate cell fate and proliferation, and canonical Wnt’s mechanism of action and biological functions are relatively well understood. In contrast, non-canonical Wnt signaling, which regulates tissue morphogenetic processes in a β-catenin-independent manner, remains poorly characterized (Clevers & Nusse, 2012; Nusse & Varmus, 2012; Steinhart & Angers, 2018; Veeman, Axelrod, & Moon, 2003). Numerous studies in a variety of model organisms have demonstrated that alterations to the expression of Wnt5a, the prototypic non-canonical Wnt ligand, can cause drastic morphogenesis defects such as body axis truncations, shortened limbs and tails, and craniofacial malformations (Hikasa, Shibata, Hiratani, & Taira, 2002; Moon et al., 1993; Yamaguchi, Bradley, McMahon, & Jones, 1999). These phenotypic abnormalities closely mirror those of Ror1 and Ror2 double knockout mice, further underscoring the growing evidence that Ror receptors mediate Wnt5a signals to orchestrate tissue morphogenetic events (Ho et al., 2012; Nomi et al., 2001).

Importantly, the phenotypic characteristics observed in *WNT5A* and *ROR1; ROR2* double mutants, namely body axis and limb truncations plus craniofacial malformations, have also been observed in human Robinow syndrome patients, and several recent publications have reported that many Robinow syndrome patients possess mutations in various components of the Wnt5a-Ror signaling pathway, including *WNT5A, ROR2, FRIZZLED2* (*FZD2*), *DISHEVELLED1* (*DVL1*), and *DISHEVELLED3* (*DVL3*) (Afzal & Jeffery, 2003; Afzal et al., 2000; Bunn et al., 2015; Person et al., 2010; J. White et al., 2015; J. J. White et al., 2018; J. J. White et al., 2016). Further, bulldogs and other closely related dog breeds possess a mutation in *DISHEVELLED2* (*DVL2*) that is highly analogous to the human Robinow syndrome mutations in *DVL1* and *DVL3*, and these breeds exhibit skeletal and craniofacial features that are reminiscent of human Robinow syndrome (Mansour et al., 2018). Collectively, these recent findings strongly support the idea that Wnt5a-Ror signaling is conserved and critical to tissue morphogenesis in a variety of vertebrates. However, despite the significance of Wnt5a-Ror signaling in both normal development and disease contexts, the mechanisms by which Wnt5a signals are transmitted and processed within the cell remain unclear. Progress within the field is further hampered by a lack of consensus regarding the number of non-canonical pathways, the biochemical nature of their regulation, and variability in the methods used to measure signaling (Veeman et al., 2003).

To deepen our understanding of Wnt5a-Ror signaling, we have taken systematic approaches to identify downstream cellular events that occur in response to pathway activation. In a previous study, we genetically ablated Ror1 and Ror2 receptors in primary mouse embryonic fibroblast (MEF) cultures and used a proteomic approach to uncover downstream signaling events that are misregulated in these cells. From this analysis, we identified the atypical kinesin Kif26b as a downstream component of Wnt5a-Ror signaling that is targeted for degradation upon pathway activation (Susman et al., 2017).

In this follow-up study, we hypothesized that additional downstream regulatory targets likely exist, and the identification of such factors would augment our mechanistic understanding of Wnt5a-Ror signaling. To that end, we conducted a second large-scale proteomic screen to identify additional cellular proteins whose abundance and phosphorylation state are altered by *acute* stimulation of Wnt5a-Ror signaling and identified the E3 ubiquitin ligase Pdzrn3 as a downstream target. Pdzrn3 has been implicated in non-canonical Wnt signaling previously (Sewduth et al., 2014). Pdzrn3 has been shown to interact with Dvl3 and influence its intracellular trafficking, and genetic loss of *PDZRN3* in human umbilical vein endothelial cells (HUVECs) decreases their cell migration in the presence of Wnt5a conditioned medium, suggesting that Pdzrn3 functions as a promigratory factor (Sewduth et al., 2014). These findings correlate with other studies that demonstrate a role for Pdzrn3 in a variety of other morphogenetic cell behaviors, including synaptic growth and maturation, vascular morphogenesis, and neuronal positioning (Baizabal et al., 2018; Lu et al., 2007; Sewduth et al., 2014). However, despite its clear involvement in modulating cell movement and positioning, how Pdzrn3 is regulated by non-canonical Wnt signaling at a biochemical level still remains unknown. We discovered that Pdzrn3 is degraded in response to Wnt5a-Ror signaling by a mechanism that is independent of β-catenin but dependent on the ubiquitin-proteasome system (UPS). This regulation is mediated by a signaling cascade involving Frizzled (Fzd), Dishevelled (Dvl), Casein kinase 1 (CK1), and Glycogen synthase kinase 3 (GSK3), which is remarkably similar to the cascade used to regulate Kif26b. Despite these similarities, we find that Wnt5a-induced degradation of Pdzrn3 is not dependent on Kif26b and vice versa, although some cross-talk exists between the two effectors. Further, Wnt5a-dependent degradation of Pdzrn3 requires phosphorylation of three specific amino acid residues on its C-terminal LNX3H domain. Critically, the phosphorylation and degradation of Pdzrn3 serves as a mechanism through which Wnt5a-Ror signaling can regulate cell migration. Lastly, we demonstrated that the LNX3H domain is required for Wnt5a-dependent degradation of not only Pdzrn3 but also its structural homolog Lnx4, suggesting that the LNX3H domain may generally function as a Wnt5a-responsive domain. Together, these findings establish the mechanisms through which the Wnt5a-Ror pathway regulates Pdzrn3 abundance to facilitate signal transduction, thus providing a platform from which a deeper mechanistic and cell morphogenetic understanding of non-canonical Wnt signaling can be attained.

## Results

### Large-scale proteomic screen identifies the E3 ubiquitin ligase Pdzrn3 as a downstream regulatory target of Wnt5a-Ror signaling

To profile both early and late molecular events driven by Wnt5a-Ror signaling, we conducted a large-scale proteomic screen in which we acutely stimulated E12.5 primary *Wnt5a* KO MEFs (Ho et al., 2012; Susman et al., 2017; Yamaguchi et al., 1999) with purified recombinant Wnt5a (rWnt5a) for 0, 1 or 6 hours, and then used quantitative tandem mass tag (TMT) mass spectrometry to globally assess changes in the abundance and phosphorylation state of cellular proteins over time (Figure 1A) (Ting, Rad, Gygi, & Haas, 2011). For rigor and reproducibility, two independent replicates of *Wnt5a* KO MEF cultures were stimulated with rWnt5a and analyzed.

**Figure 1.**
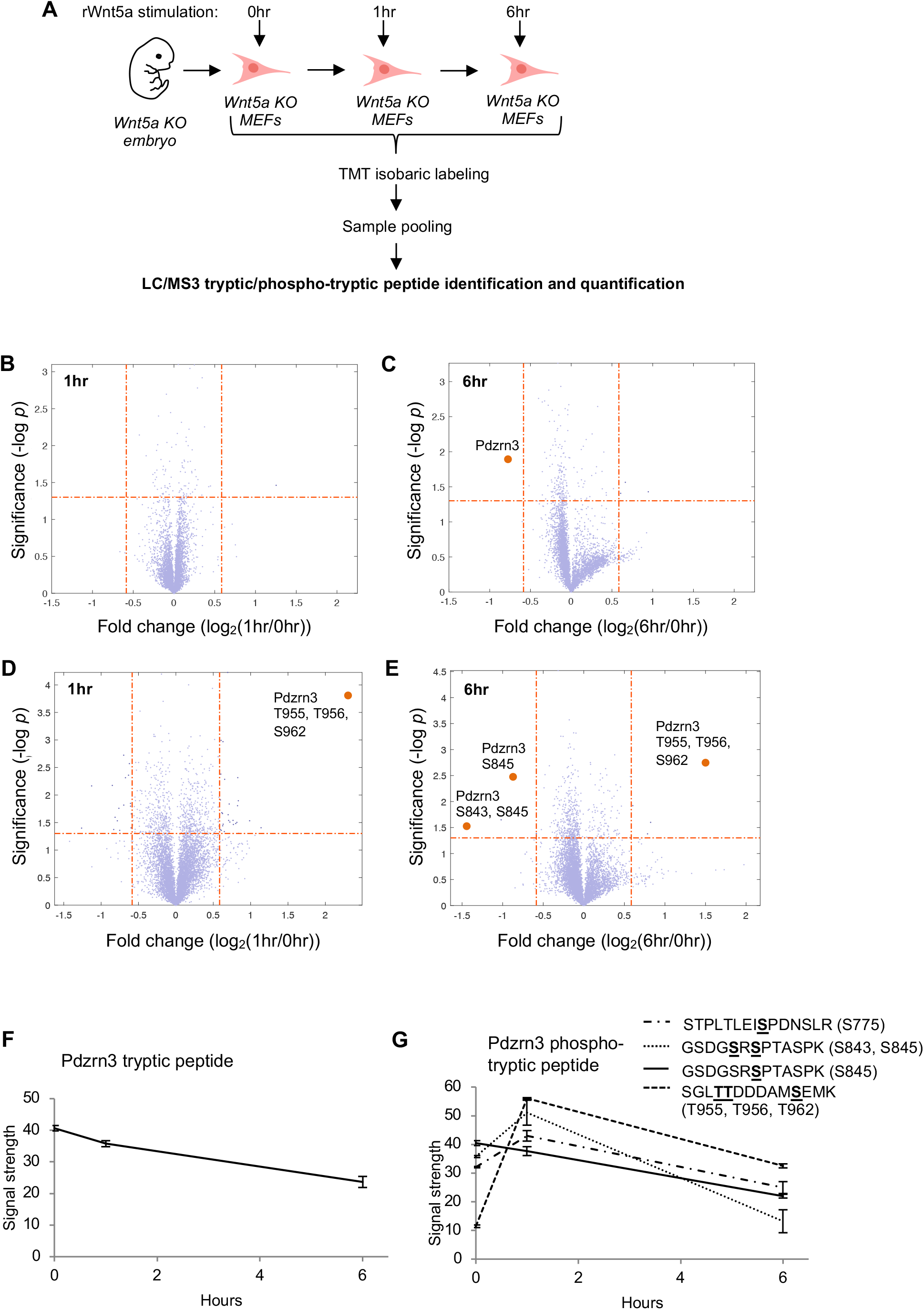
Identification of the E3 ubiquitin ligase Pdzrn3 as a downstream regulatory target of Wnt5a-Ror signaling. **A)** Workflow of whole cell proteomics screen. Primary MEF cultures were generated from *Wnt5a* knockout E12.5 mouse embryos and stimulated with rWnt5a (0.1μg/mL) for 0, 1, or 6 hours. After rWnt5a stimulation, whole cell lysates were collected and processed for LC/MS3 tryptic/phospho-tryptic peptide identification and quantification, as described in the main text and Materials and Methods. The rWnt5a stimulation and proteomic analysis were conducted in two independent technical replicates. **B and C**) Volcano plots showing changes in the abundance of detected tryptic peptides in response to rWnt5a stimulation (0.1μg/mL) after 1 hour (B) or 6 hours (C). The abundance of a tryptic peptide from Pdzrn3 (orange dots) changed strongly after 6 hours of rWnt5a stimulation. **D and E)** Volcano plots showing changes in the abundance of detected phospho-tryptic peptides after 1 hour (D) or 6 hours (E) of rWnt5a stimulation (0.1μg/mL). A total of five phosphosites from Pdzrn3 (S843, S845, T955, T956, and S962), grouped in two clusters based on their location in the protein, were detected and exhibited distinct patterns of change after rWnt5a stimulation (orange dots). **F and G)** Line plots showing Wnt5a-induced changes in the abundance of individual tryptic (F) or phospho-tryptic peptides (G) from Pdzrn3 after 1 hour or 6 hours of rWnt5a stimulation. A sixth phosphor-tryptic peptide site (S775) in did not pass the initial filter (p-value = 0.076), but also showed clear changes with rWnt5a stimulation. Error bars represent ± SEM calculated from two technical replicates.

In both analyses of protein level and phosphorylation changes, we defined potential proteins of interest as those with tryptic peptides or phospho-tryptic peptides that exhibited (1) a negative or positive change of > 1.5-fold in abundance and (2) a change with a p-value < 0.05 across the two experimental replicates. Based on these criteria, our top candidate was Pdzrn3, an E3 ubiquitin ligase, which exhibited significant changes in both steady-state protein abundance and phosphorylation state after rWnt5a stimulation. Although 1 hour of rWnt5a stimulation did not result in detectable changes in Pdzrn3 abundance (Figure 1B), after 6 hours of rWnt5a stimulation we observed that Pdzrn3 abundance was significantly downregulated by 1.72-fold (p= 0.013, Figure 1C and 1F; Table 1). Additionally, we identified multiple phospho-tryptic peptides derived from two different regions of Pdzrn3 that exhibited significant changes after rWnt5a stimulation (Figure 1D and 1E). A phospho-tryptic peptide containing S843 and S845 (dotted line, Figure 1G) and another one containing T955, T956 and S962 (dashed line, Figure 1G) both showed an initial increase at 1 hour, followed by a decrease at 6 hours. Likewise, a third phospho-tryptic peptide containing S775 (dot dashed line, Figure 1G), though not scored initially based on its significance value (p=0.076), also exhibited a similar pattern of change. Lastly, a phospho-tryptic peptide containing S845 (solid line, Figure 1G) decreased gradually after 1 hour and more extensively after 6 hours. Importantly, all phospho-tryptic peptides decrease by 6 hours to a similar extent as that of the non-phosphorylated tryptic peptide (Figure 1F). This overall pattern thus raised the hypothesis that Wnt5a signaling first induces the phosphorylation of Pdzrn3 at specific sites at 1 hour, followed by downregulation of Pdzrn3 protein abundance at 6 hours, and these two biochemical events are kinetically and mechanistically coupled.

**Table 1.**
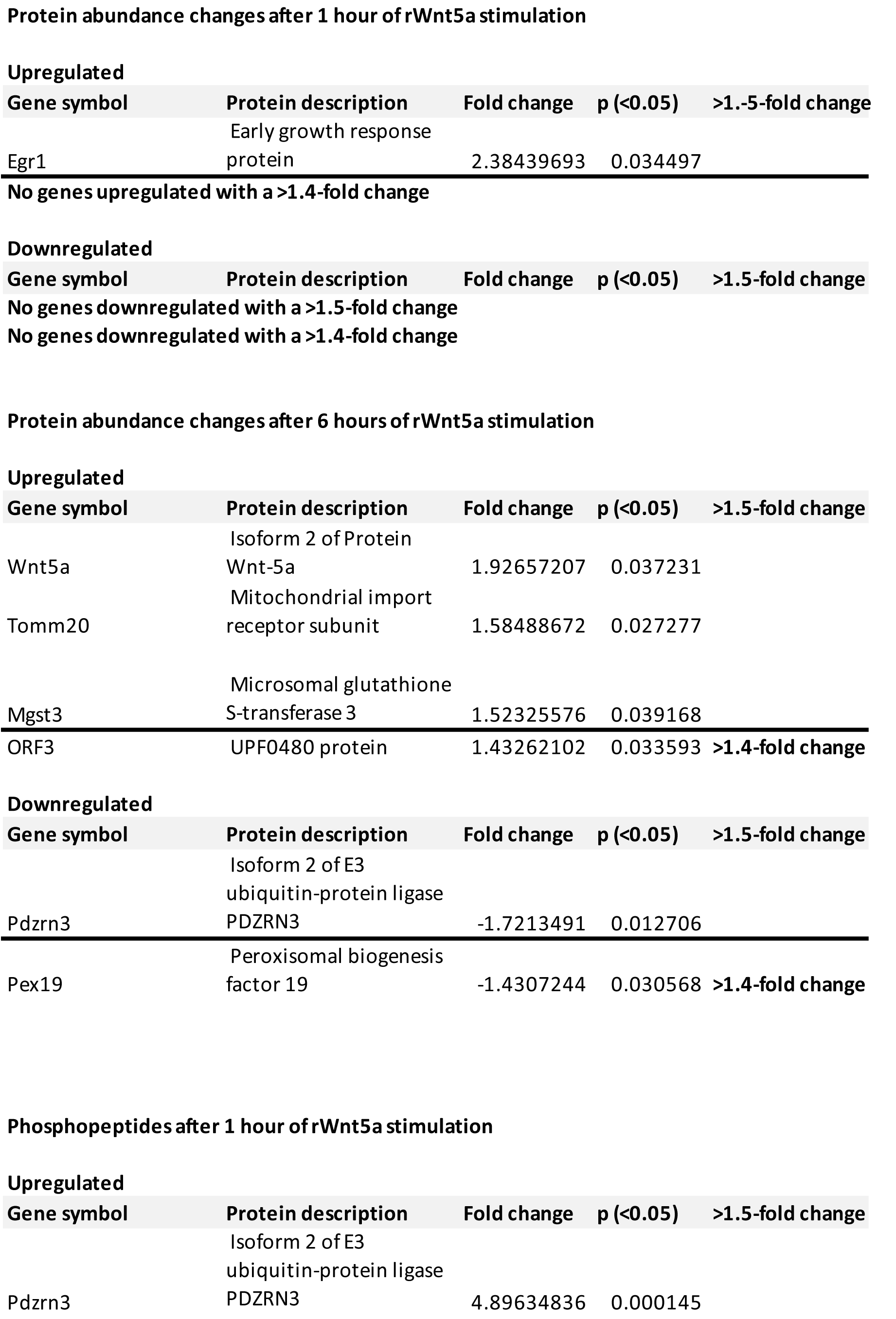
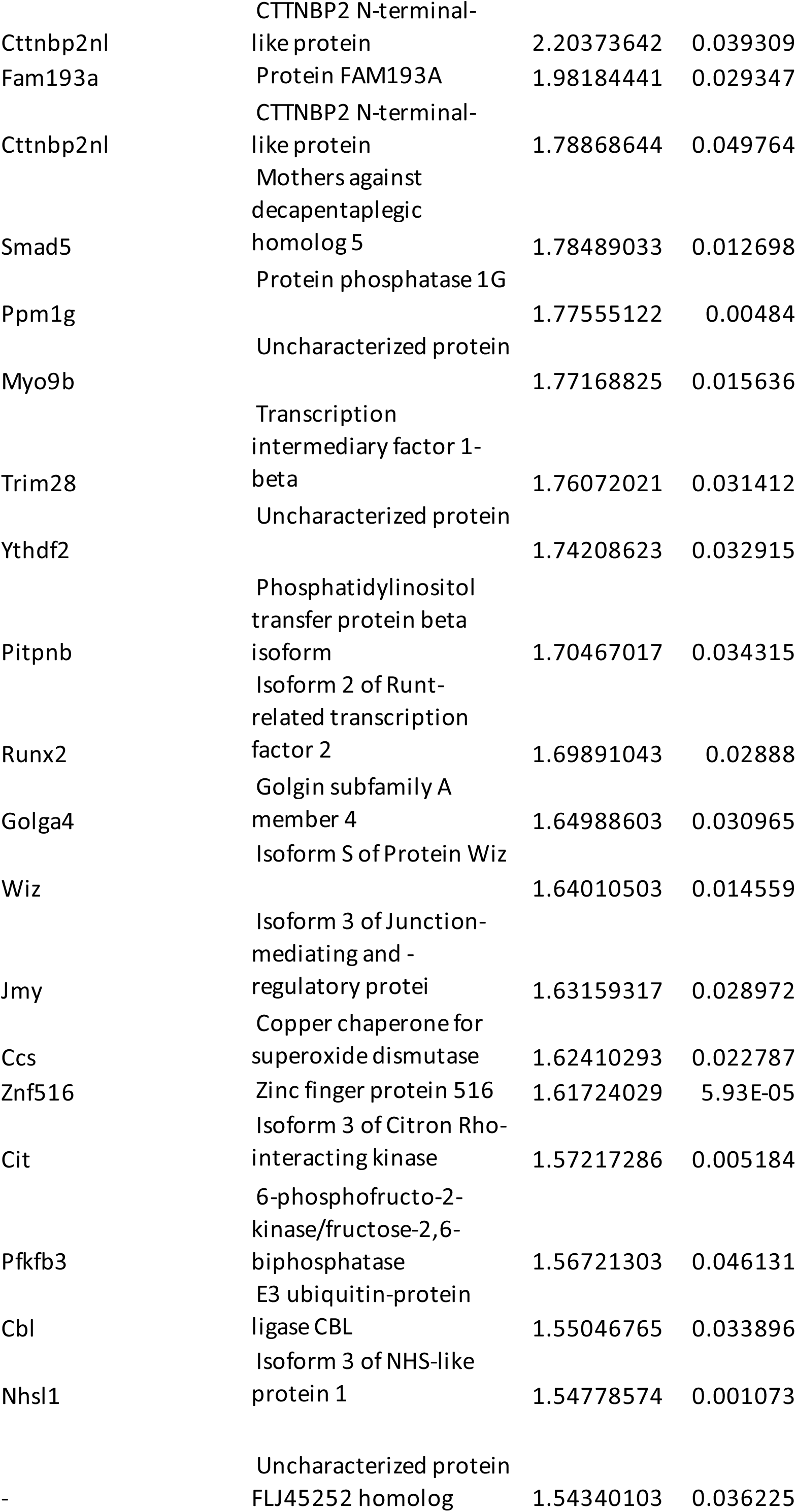
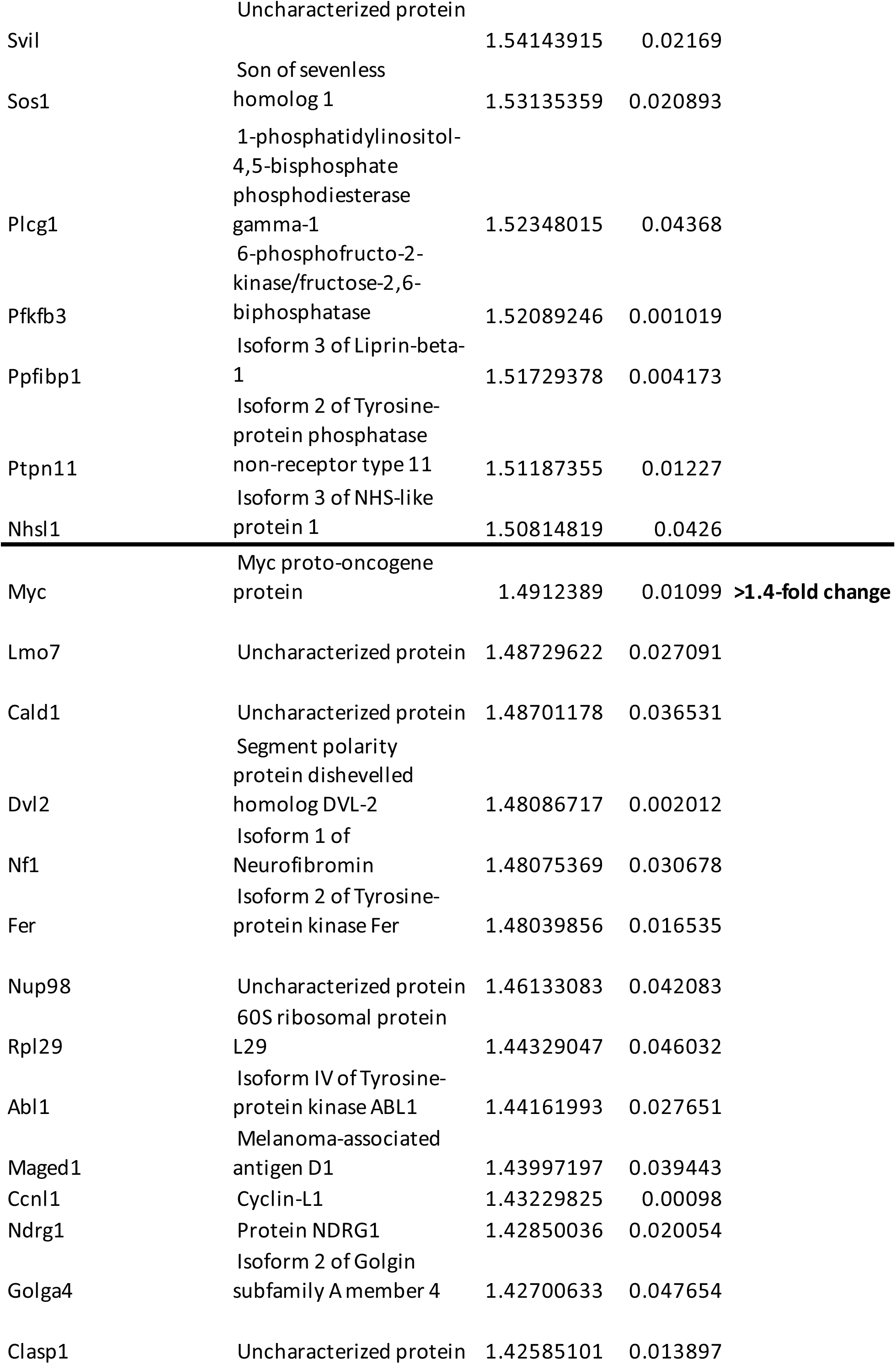
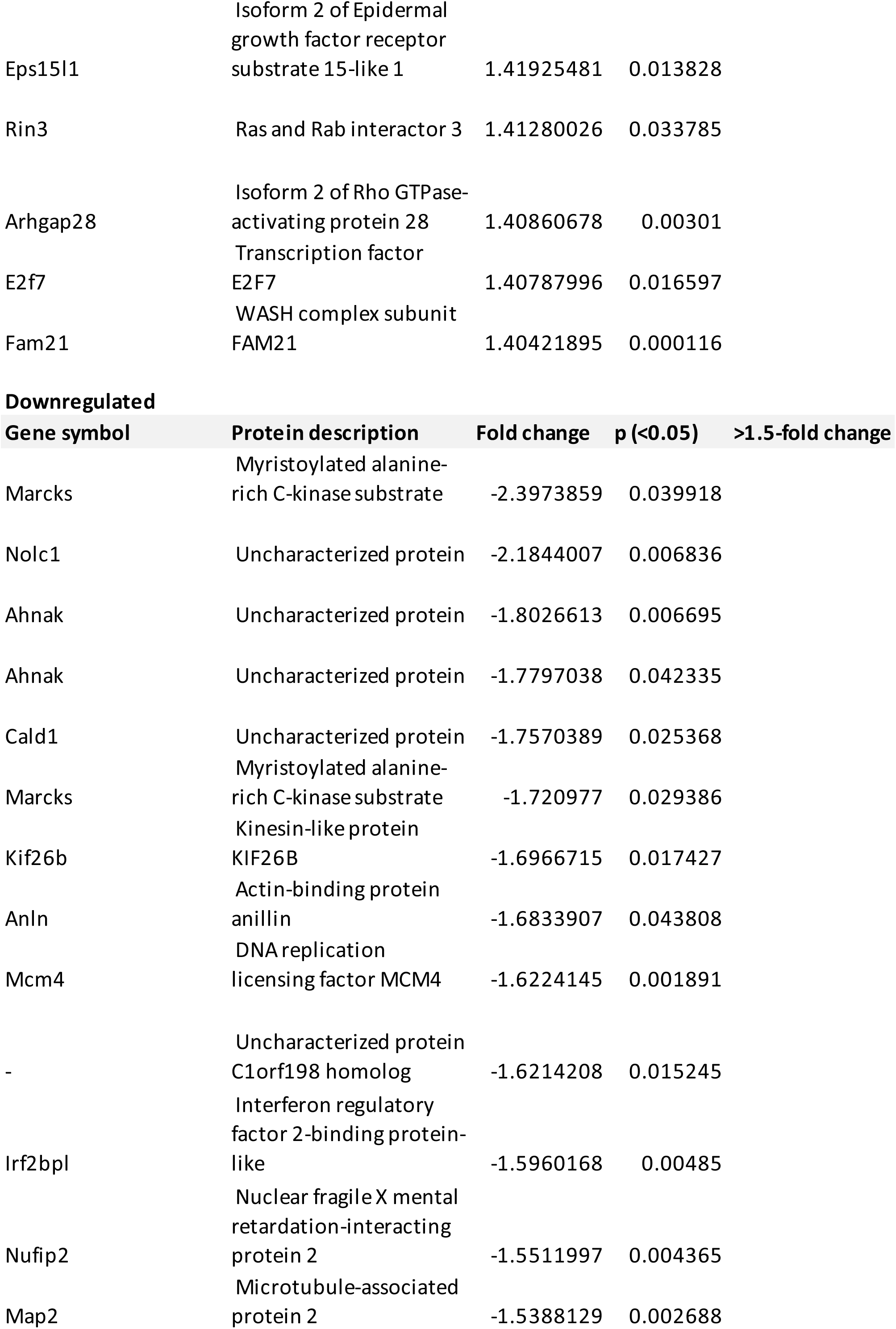
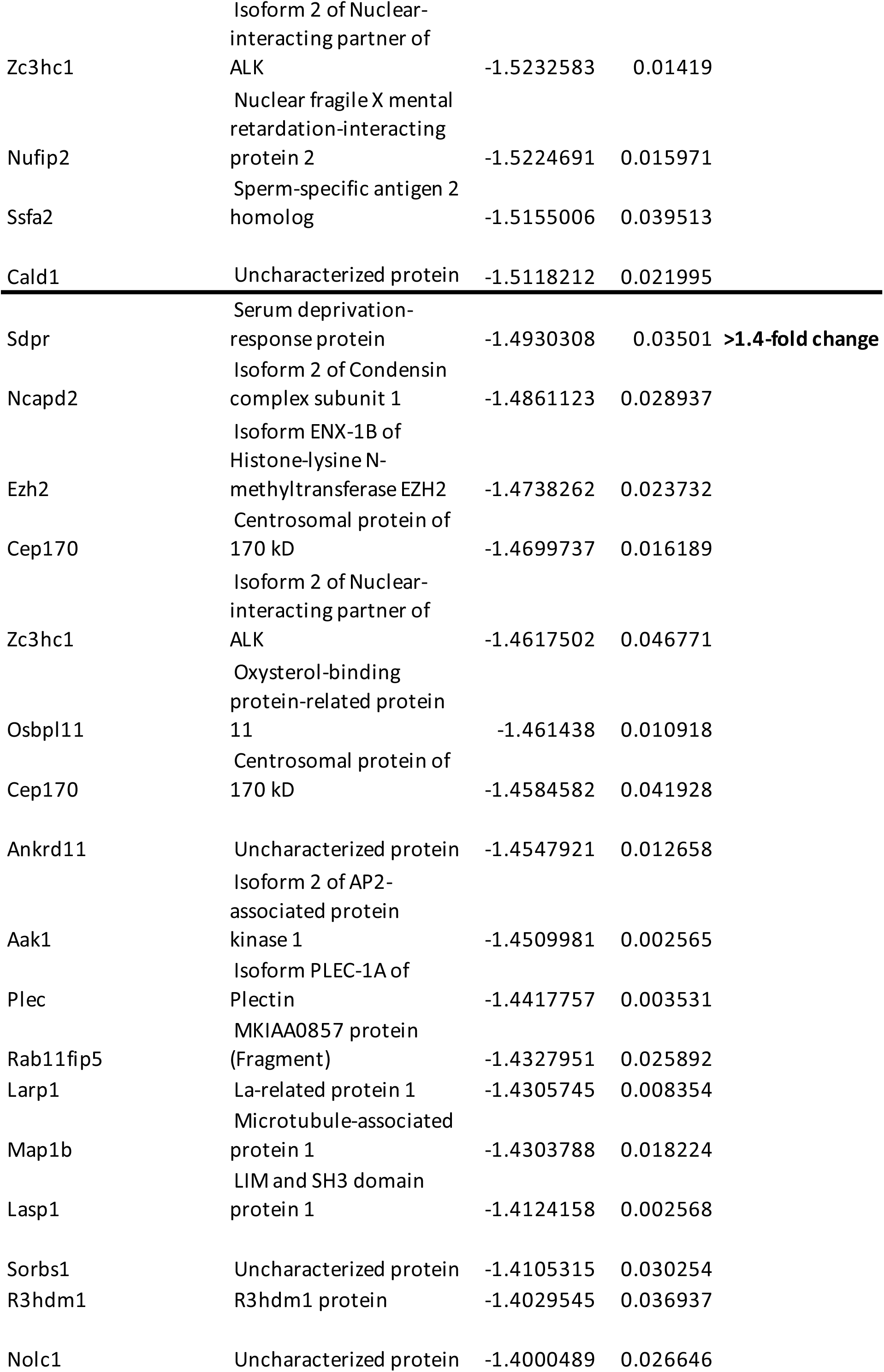
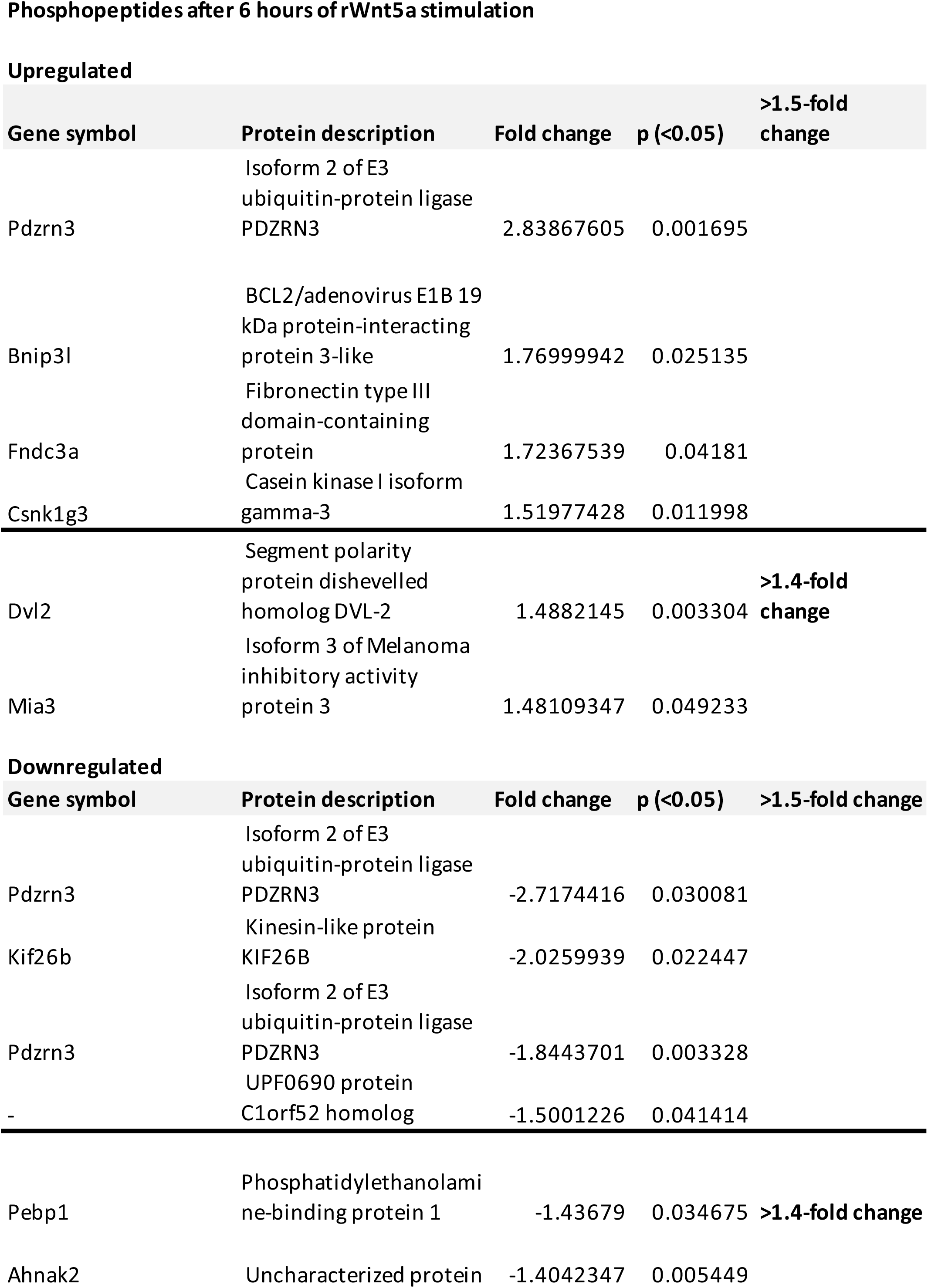
Hits from Wnt5a knockout MEF TMT/MS3 screen.

In addition to Pdzrn3, the proteomic screen also identified several other known components of the Wnt5a-Ror signaling pathway (Table 1). At both the 1 hour and 6 hour timepoints, a phospho-tryptic peptide from Kif26b was scored as a “hit” (1.70- and 2.03-fold decrease, respectively). At the 6 hour timepoint, a phospho-tryptic peptide from CK1 isoform gamma-3 was also scored as a “hit” (1.52-fold increase). In addition, a phospho-tryptic peptide from Dvl2 exhibited a 1.49-fold increase in abundance after 6 hours of rWnt5a stimulation. The identification of these previously described Wnt5a signaling targets further validates the selectivity and sensitivity of the proteomic screening approach.

To independently confirm that Pdzrn3 abundance is indeed regulated by Wnt5a signals, we generated rabbit polyclonal antibodies against Pdzrn3 and used western blotting to analyze the steady-state cellular levels of Pdzrn3 after rWnt5a stimulation. Consistent with our proteomic screening results, we observed that the abundance of Pdzrn3 significantly decreased after 6 hours of rWnt5a stimulation (Figure 2A and 2B). This change parallels other previously described responses of Wnt5a-Ror signaling, including an increase in the phosphorylation of Ror1, Ror2 and Dvl2, and a decrease in Kif26b abundance (Figure 2A and 2B) (Ho et al., 2012; Susman et al., 2017).

**Figure 2.**
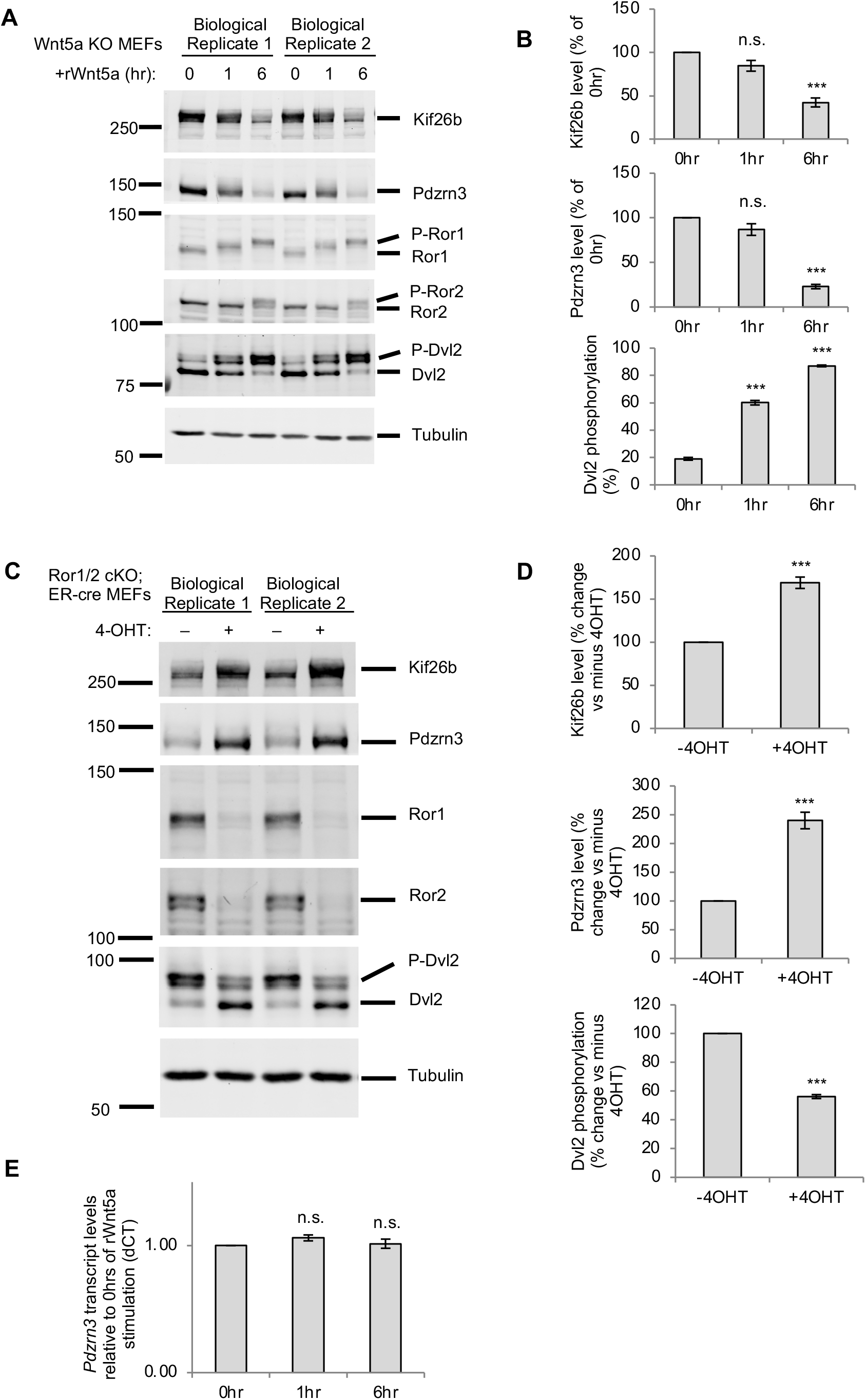
Validation of Pdzrn3 as a downstream regulatory target of Wnt5a-Ror signaling. **A)** Western blot showing downregulation of Pdzrn3 steady-state levels in response to rWnt5a stimulation. Primary *Wnt5a* knockout MEF cultures (n=3, biological replicates) were stimulated with rWnt5a (0.2μg/mL) for 0, 1, or 6 hours, and membranes were blotted with antibodies against Kif26b, Pdzrn3, Ror1, Ror2, Dvl2 and Tubulin. The decrease in Pdzrn3 abundance correlated with other known indicators of Wnt5a-Ror signaling, including phosphorylation changes in Ror1, Ror2, and Dvl2 and a decrease in Kif26b abundance. Experiments from two representative biological replicates are shown. **B)** Quantification of the western blotting experiments shown in (A). Error bars represent ± SEM calculated from three biological replicates. t-test (unpaired) was performed to determine statistical significance for the following comparisons: 1hr vs. 0hr; 6hr vs. 0hr. **C)** Western blots showing the requirement of endogenous Ror receptors for Pdzrn3 regulation. Primary MEFs derived from *Ror1*^*fl/fl*^; *Ror2*^*fl/fl*^; *ER-cre* embryos were treated with 4-hydroxytamoxifen (4-OHT) to induce genetic ablation of *Ror1* and *Ror2*. Protein lysates were analyzed by western blotting using antibodies against Kif26b, Pdzrn3, Ror1, Ror2, Dvl2 and Tubulin. An increase in Pdzrn3 and Kif26b steady-state abundance along with a decrease in Dvl2 phosphorylation correlated with the genetic loss of Ror1 and Ror2 expression. **D)** Quantification of the western blotting experiments shown in (C). Error bars represent ± SEM calculated from three biological replicates. t-test (unpaired) was performed to determine statistical significance for the following comparisons: +4OHT vs. −4OHT. **E)** Plot showing the effect of Wnt5a stimulation on *Pdzrn3* transcript levels. Primary *Wnt5a* knockout MEFs were stimulated with rWnt5a (0.2μg/mL) for 0, 1, or 6 hours, and the relative abundance of *Pdzrn3* mRNA were determine by RT-qPCR. Error bars represent ± SEM calculated from three technical replicates. t-test (unpaired) was performed to determine statistical significance for the following comparisons: 1hr vs. 0hr; 6hr vs. 0hr. P-values: * = p<0.05, ** = p<0.01, *** = p<0.001.

To test whether Ror receptors are required for Wnt5a signaling to Pdzrn3, we took advantage of conditional Ror receptor family knockout MEFs derived from E12.5 *Ror1*^*fl/fl*^; *Ror2*^*fl/fl*^; *CAG-CreER* embryos. These MEFs undergo robust autocrine/paracrine Wnt5a-Ror signaling even in the absence of exogenously added Wnt5a (Ho et al., 2012). To genetically ablate *Ror1* and *Ror2* expression in these MEFs, we treated the cells with 4-hydroxytamoxifen (4OHT) to induce CreER-mediated deletion of the *Ror1*^*fl/fl*^ and *Ror2*^*fl/fl*^ alleles. We observed that loss of Ror receptor expression resulted in a significant increase in Pdzrn3 levels, which correlated with a decrease in Dvl2 phosphorylation as well as an increase in Kif26b abundance (Figure 2C and 2D). Thus, these results indicate that, indeed, Ror receptors are required to facilitate Wnt5a-driven regulation of Pdzrn3 abundance, and that this regulation is a genuine endogenous Wnt5a-Ror signaling event.

To test whether Wnt5a regulation of Pdzrn3 protein abundance occurs transcriptionally or post-transcriptionally, we treated *Wnt5a* knockout MEFs with rWnt5a for 1 or 6 hours and analyzed the levels of *Pdzrn3* mRNA by reverse transcription-quantitative polymerase chain reaction (RT-qPCR). Unlike Pdzrn3 protein, *Pdzrn3* transcripts do not change significantly after 1 or 6 hours of rWnt5a stimulation (Figure 2E). Thus, Wnt5a-Ror signaling regulates Pdzrn3 protein abundance through a post-transcriptional mechanism. Overall, these experiments establish that regulation of Pdzrn3 protein abundance is a physiological response of Wnt5a-Ror signaling.

### A non-canonical Wnt signaling cascade involving Fzd, Dvl, CK1, GSK3 and the ubiquitin-proteasome system regulates Pdzrn3 degradation

To dissect the molecular mechanisms that mediate Wnt5a regulation of Pdzrn3, we designed a flow cytometry-based reporter in which we stably expressed GFP-Pdzrn3 in NIH/3T3 cells (referred to as WRP reporter cells for <u>W</u>nt5a-<u>R</u>or-<u>P</u>dzrn3). Consistent with our observations in primary *Wnt5a* knockout MEFs, treatment of the WRP reporter cells with rWnt5a for 6 hours, under conditions in which endogenous Wnt signaling is inhibited with the small molecule PORCN inhibitor Wnt-C59, resulted in a significant downregulation of GFP-Pdzrn3 reporter signal, thereby demonstrating the fidelity of this reporter assay (Figure 3A). Moreover, we established that a saturable dose-dependent relationship exists between rWnt5a concentrations and GFP-Pdzrn3 downregulation, with a calculated EC_50_ of 77.1 ng/mL, which is similar to other Wnt induced responses (Figure 3B) (Bryja, Schulte, Rawal, Grahn, & Arenas, 2007; Connacher, Tay, & Ahn, 2017; Ho et al., 2012; Park et al., 2015; Witze et al., 2013; Witze, Litman, Argast, Moon, & Ahn, 2008). This supports the physiological relevance of Pdzrn3 downregulation.

**Figure 3.**
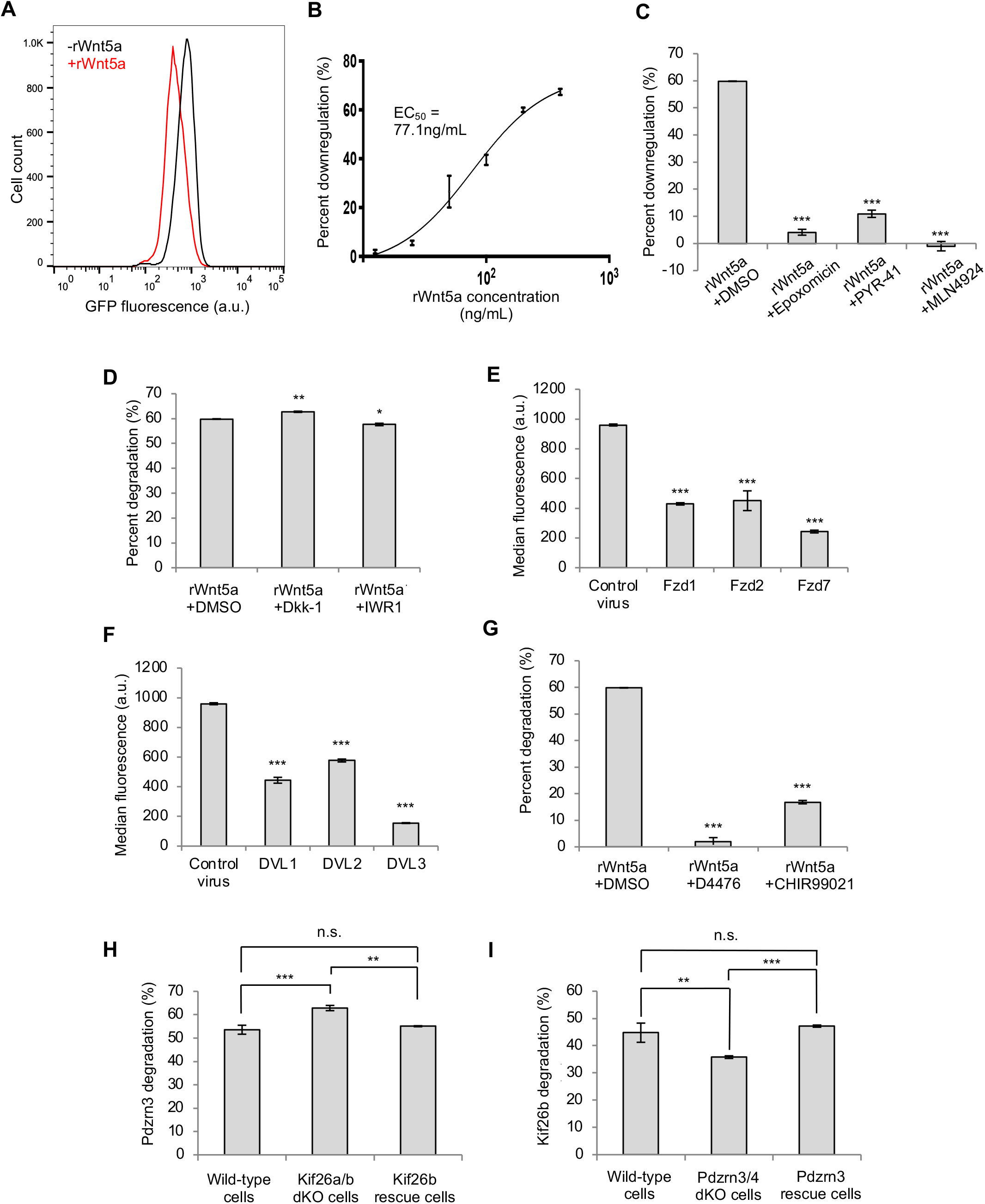
A signaling cascade links non-canonical Wnt5a-Ror signaling to Pdzrn3 degradation. **A)** Representative histogram showing the effect of rWnt5a treatment on NIH/3T3 GFP-Pdzrn3 (WRP) reporter cells. WRP cells were treated with rWnt5a (0.2μg/mL) for 6 hours, and GFP-Pdzrn3 fluorescence was measured by flow cytometry. **B)** Dose-response curve showing GFP-Pdzrn3 downregulation as a function of rWnt5a concentration in the WRP reporter assay. An EC_50_ of 77.1ng/mL was calculated. **C)** Quantification of the effects of proteasome inhibitor (epoxomicin, 10μM), ubiquitin-activating enzyme E1 inhibitor (PYR41, 10μM) and Cullin inhibitor (MLN4924, 10μM) on rWnt5a-induced Pdzrn3 downregulation in the WRP reporter cells. Error bars represent ± SEM calculated from three technical replicates. t-test (unpaired) was performed to determine statistical significance for the following comparisons: inhibitors vs. the vehicle control DMSO. **D)** Quantification of the effects of canonical Wnt inhibitors, Dkk-1 (2μg/μL) and IWR-1-endo (10μM) on rWnt5a-induced Pdzrn3 degradation in the WRP reporter cells. Error bars represent ± SEM calculated from three technical replicates. t-test (unpaired) was performed to determine statistical significance for the following comparisons: inhibitors vs. the vehicle control DMSO. **E)** Quantification of the effects of Fzd1, Fzd2 and Fzd7 overexpression on the median fluorescence of WRP reporter cells. Error bars represent ± SEM calculated from two cell lines and three technical replicates per line. t-test (unpaired) was performed to determine statistical significance for the following comparisons: Fzd overexpression vs. the Myc epitope tag overexpression. **F)** Quantification of the effects of DVL1, DVL2 and DVL3 overexpression on the median fluorescence of WRP reporter cells. Error bars represent ± SEM calculated from two cell lines and three technical replicates per line. t-test (unpaired) was performed to determine statistical significance for the following comparisons: DVL overexpression vs. the Myc epitope tag overexpression. **G)** Quantification of the effects of CK1 inhibitor (D4476, 100μM) and GSK inhibitor (CHIR99021, 100μM) on rWnt5a-induced Pdzrn3 downregulation in the WRP reporter cells. Error bars represent ± SEM calculated from three technical replicates. t-test (unpaired) was performed to determine statistical significance for the following comparisons: inhibitors vs. the vehicle control DMSO. **H)** Quantification of the effect of genetically ablating *Kif26a* and *Kif26b* on rWnt5a-induced GFP-Pdzrn3 reporter degradation. Error bars represent ± SEM calculated from three technical replicates. t-test (unpaired) was performed to determine statistical significance for the following comparisons: GFP-Pdzrn3 reporter in *Kif26a/Kif26b* double KO cells vs. GFP-Pdzrn3 reporter in WT cells; GFP-Pdzrn3 reporter in Kif26b rescue cells vs. GFP-Pdzrn3 reporter in WT cells; GFP-Pdzrn3 reporter in *Kif26a/Kif26b* double KO cells vs. GFP-Pdzrn3 reporter in Kif26b rescue cells. **I)** Quantification of the effect of genetically ablating *Pdzrn3* and *Lnx4* on rWnt5a-induced GFP-Kif26b reporter (WRK) degradation. Error bars represent ± SEM calculated from three technical replicates per line. t-test (unpaired) was performed to determine statistical significance for the following comparisons: GFP-Kif26b reporter in *Pdzrn3/Lnx4* double KO cells vs. GFP-Kif26b reporter in WT cells; GFP-Kif26b reporter in Pdzrn3 rescue cells vs. GFP-Kif26b reporter in WT cells; GFP-Kif26b reporter in *Pdzrn3/Lnx4* double KO cells vs. GFP-Kif26b reporter in Pdzrn3 rescue cells. P-values: * = p<0.05, ** = p<0.01, *** = p<0.001.

**Figure 3.**
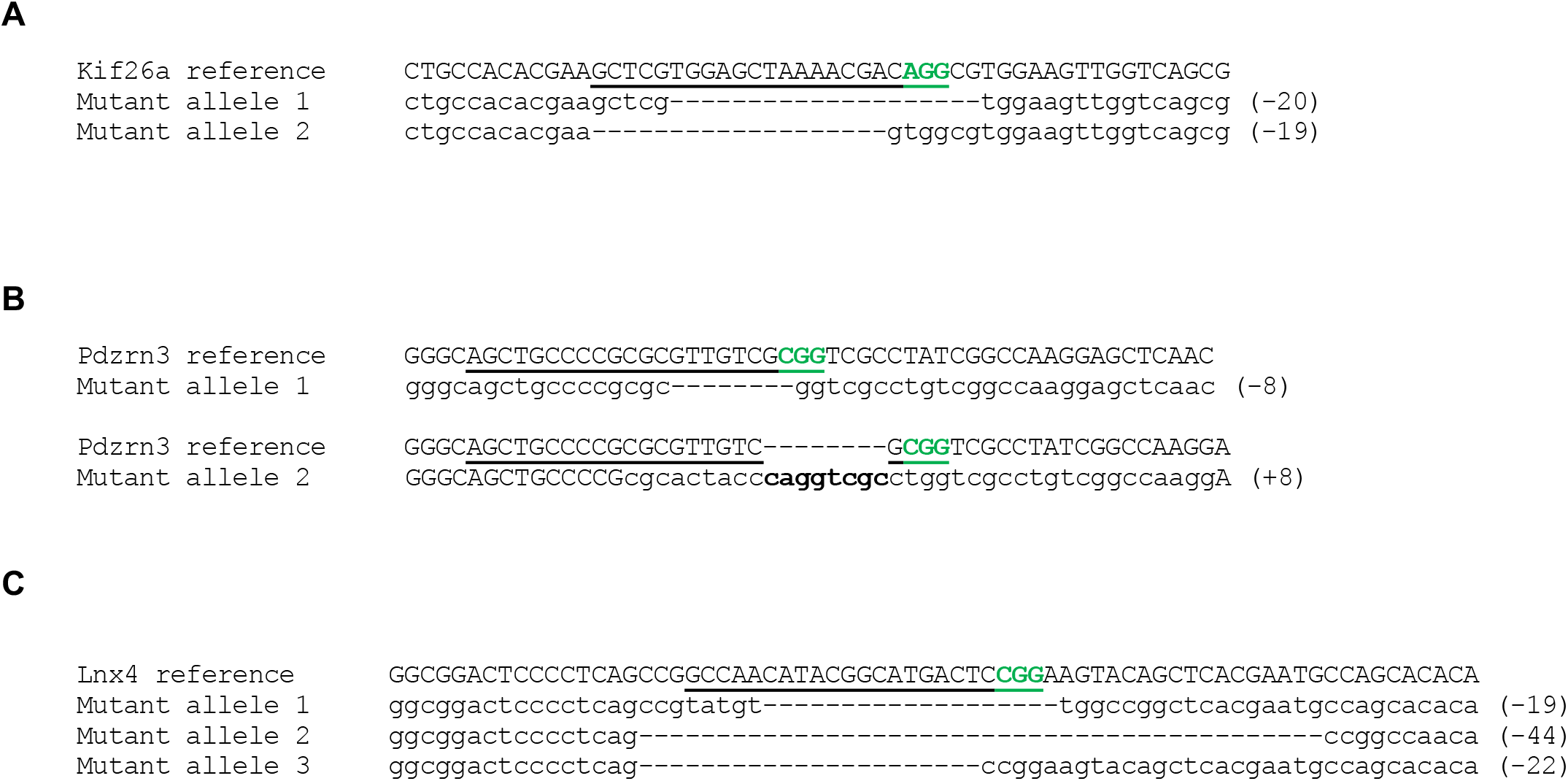
Supplement. CRISPR/Cas9 mediated genetic deletions of *Kif26a, Pdzrn3*, and *Lnx4*. Reference sequences for *Kif26a* (A), *Pdzrn3* (B), and *Lnx4* (C) aligned to mutant alleles generated via targeting with short guide RNAs (sgRNAs) (underlined in reference, green refers to PAM sequence) unique to each gene. Multiple deep sequencing results indicate that *Lnx4* is triploid, which is consistent with previous karyotyping of NIH/3T3 cells (Leibiger et al., 2013).

**Figure 4.**
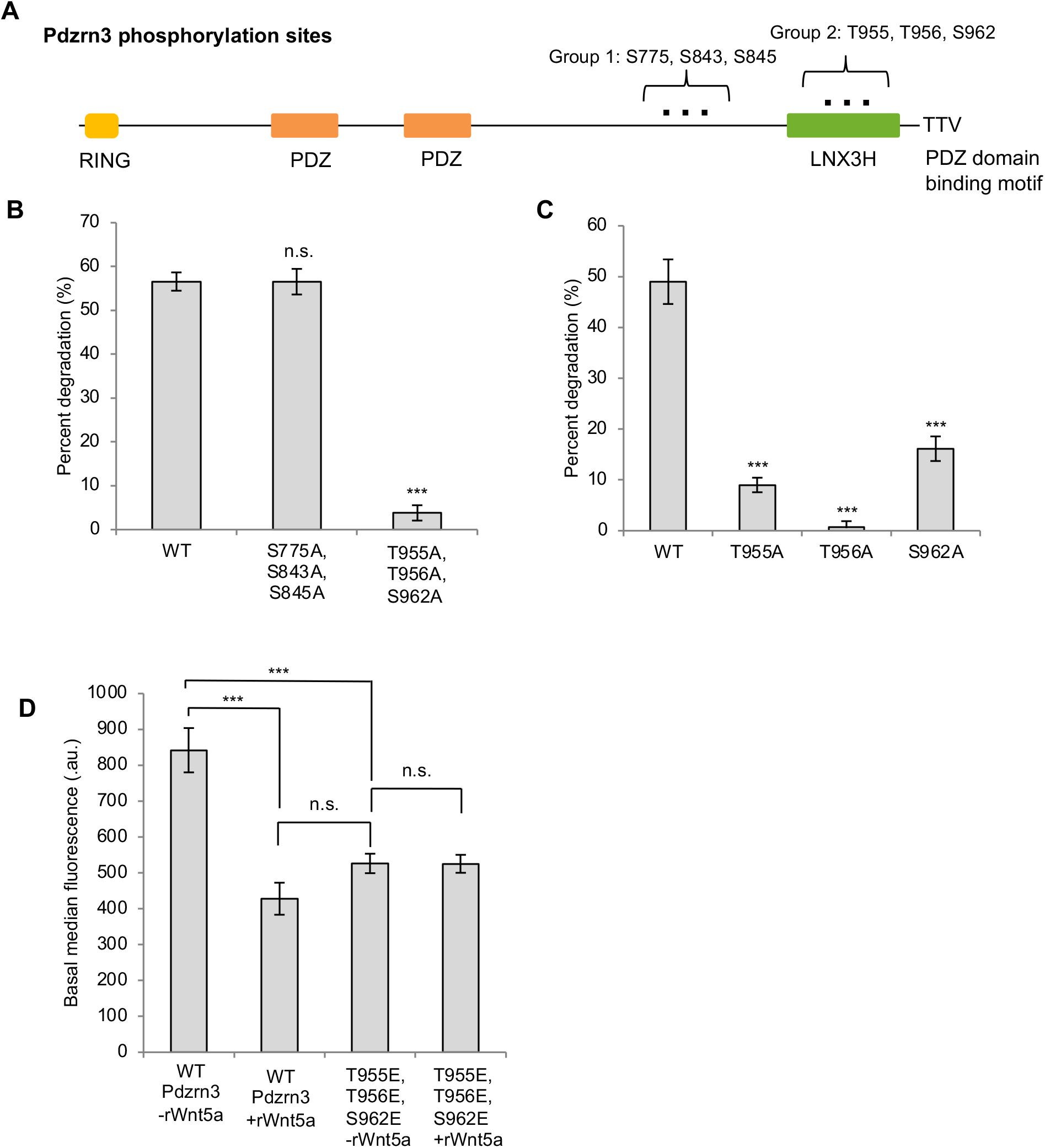
Pdzrn3 phosphorylation is required for its degradation. **A)** Schematic of domains and identified phosphorylation sites of Pdzrn3. **B)** Quantification of the effects of mutating Group 1 and Group 2 sites on Wnt5a-induced Pdzrn3 degradation. Error bars represent ± SEM calculated from two cell lines and three technical replicates per line. t-test (unpaired) was performed to determine statistical significance for the following comparisons: mutant Pdzrn3 vs. WT Pdzrn3. **C)** Quantification of the effects of mutating individual Group 2 sites on Wnt5a-induced Pdzrn3 degradation. Error bars represent ± SEM calculated from two cell lines and three technical replicates per line. t-test (unpaired) was performed to determine statistical significance for the following comparisons: mutant Pdzrn3 vs. WT Pdzrn3. **D)** Quantification of the effects of Group 2 phosphomimetic mutations on Pdzrn3 reporter signals. Error bars represent ± SEM calculated from two cell lines and three technical replicates per line. t-test (unpaired) was performed to determine statistical significance for the following comparisons: WT Pdzrn3 – rWnt5a vs. WT Pdzrn3 + rWnt5a; phosphomimetic Pdzrn3 mutant – rWnt5a vs. WT Pdzrn3 -rWnt5a; phosphomimetic Pdzrn3 mutant -rWnt5a vs. WT Pdzrn3 +rWnt5a; phosphomimetic Pdzrn3 mutant + rWnt5a vs. –rWnt5a. P-values: * = p<0.05, ** = p<0.01, *** = p<0.001.

**Figure 5.**
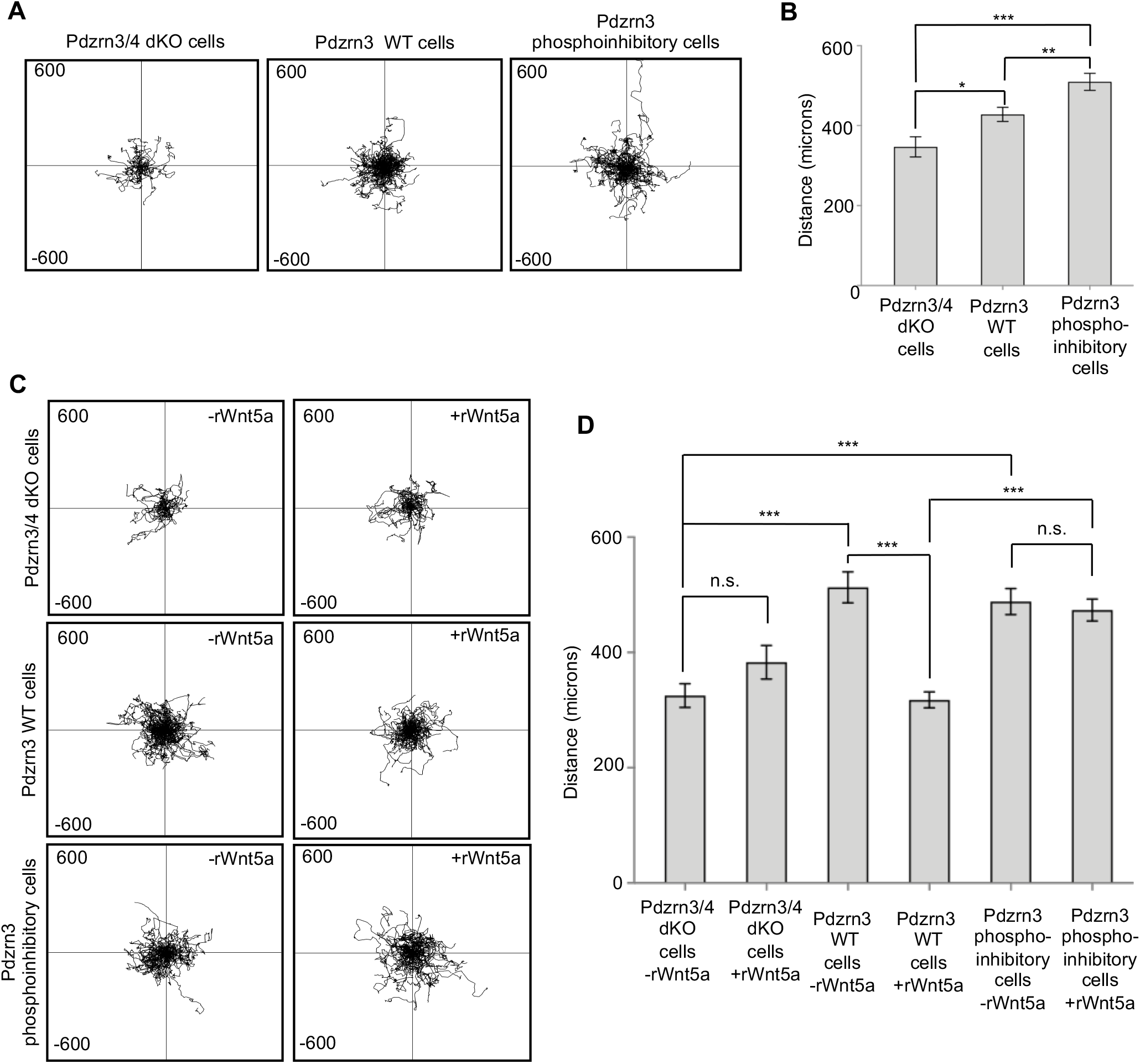
Wnt5a-dependent cell migration is regulated through Pdzrn3 phosphorylation and degradation. **A)** Single cell tracking plots of Pdzrn3 and Lnx4 double knockout cells (Pdzrn3/4 dKO cells), Pdzrn3/4 dKO cells re-expressing wild-type Pdzrn3 (Pdzrn3 WT cells), or Pdzrn3/4 dKO cells re-expressing Pdzrn3 with phosphoinhibitory mutations at Group 2 sites (Pdzrn3 phosphoinhibitory cells) without any Wnt-C59 or rWnt5a treatments. X and Y axes extend to 600 microns. **B)** Quantification of distance traversed during cell migration by Pdzrn3/4 dKO cells, Pdzrn3 WT cells, and Pdzrn3 phosphoinhibitory cells. Error bars represent ± SEM calculated from one (Pdzrn3/4 dKO cells) or two (Pdzrn3 WT cells and Pdzrn3 phosphoinhibitory cells) independent cell lines and two technical replicates per line. t-test (unpaired) was performed via Prism 8 (GraphPad Software) to determine statistical significance for the following comparisons: Pdzrn3/4 dKO cells vs. Pdzrn3 WT cells; Pdzrn3/4 dKO cells vs. Pdzrn3 phosphoinhibitory cells; Pdzrn3 WT cells vs. Pdzrn3 phosphoinhibitory cells. **C)** Single cell tracking plots of Pdzrn3/4 dKO cells, Pdzrn3 WT cells, and Pdzrn3 phosphoinhibitory cells treated with or without rWnt5a in the presence of Wnt-C59. X and Y axes extend to 600 microns. **D)** Quantification of distance traversed during cell migration by Pdzrn3/4 dKO cells, Pdzrn3 WT cells, or Pdzrn3 phosphoinhibitory cells treated with or without rWnt5a. Error bars represent ± SEM calculated from one (Pdzrn3/4 dKO cells) or two independent (Pdzrn3 WT cells and Pdzrn3 phosphoinhibitory cells) cell lines and two technical replicates per line. t-test (unpaired) was performed via Prism 8 (GraphPad Software) to determine statistical significance for the following comparisons: Pdzrn3/4 dKO cells -rWnt5a vs. Pdzrn3/4 dKO cells +rWnt5a; Pdzrn3 WT cells -rWnt5a vs. Pdzrn3 WT cells +rWnt5a; Pdzrn3 phosphoinhibitory cells -rWnt5a vs. Pdzrn3 phosphoinhibitory cells +rWnt5a; Pdzrn3/4 dKO cells -rWnt5a vs. Pdzrn3 WT cells -rWnt5a; Pdzrn3/4 dKO cells -rWnt5a vs. Pdzrn3 phosphoinhibitory cells -rWnt5a; Pdzrn3 WT cells +rWnt5a vs. Pdzrn3 phosphoinhibitory cells +rWnt5a. P-values: * = p<0.05, ** = p<0.01, *** = p<0.001.

We next used the WRP reporter cells to investigate the biochemical nature of Pdzrn3 downregulation. We pharmacologically tested the role of the ubiquitin-proteasome system (UPS) in Pdzrn3 downregulation as the UPS is a major regulatory pathway involved in many signaling systems, and our previous study demonstrated that it is required for Wnt5a-dependent degradation of Kif26b (Susman et al., 2017). We treated WRP cells with a panel of small-molecule inhibitors that block different components of the UPS: epoxomicin, which targets the proteasome (Meng et al., 1999); PYR-41, which targets the ubiquitin-activating enzyme E1 (Yang et al., 2007); and MLN4924, which targets Cullin E3 ligases (Tong et al., 2017). Each of these drugs significantly inhibited Wnt5a-dependent Pdzrn3 downregulation (Figure 3C), indicating that the UPS and the Cullin family of E3 ligases are required for Wnt5a-dependent degradation of Pdzrn3.

To test whether Wnt5a-Ror-dependent Pdzrn3 degradation occurs via a non-canonical Wnt signaling mechanism independent of the Wnt/β-catenin pathway, we treated WRP reporter cells with Dkk-1 and IWR-1-endo, which block canonical Wnt/β-catenin signaling at the receptor and destruction complex level, respectively (Bafico, Liu, Yaniv, Gazit, & Aaronson, 2001; Huang et al., 2009; E. Lee, Salic, Kruger, Heinrich, & Kirschner, 2003). We observed that neither inhibitor blocked Wnt5a-induced degradation of GFP-Pdzrn3, indicating that this regulation occurs independently of the canonical Wnt pathway (Figure 3D). However, we noted that both inhibitors, in the absence of Wnt5a treatment, slightly increased the basal fluorescence of the GPF-Pdzrn3 reporter; the mechanism behind this regulation is currently unclear. Nevertheless, our data demonstrate that Wnt5a-Ror-Pdzrn3 signaling is a bona fide non-canonical Wnt pathway.

We next investigated if other established Wnt signaling mediators are also involved in Pdzrn3 degradation. We focused our analysis on the Fzd1, Fzd2, and Fzd7 subfamily of Fzd receptors and all three members of the family of Dvl scaffolding proteins based on their emerging connection to Robinow syndrome (Afzal & Jeffery, 2003; Afzal et al., 2000; Bunn et al., 2015; Person et al., 2010; J. White et al., 2015; J. J. White et al., 2018; J. J. White et al., 2016). We overexpressed mouse Fzd1, Fzd2, or Fzd7 and human DVL1, DVL2 or DVL3 in WRP reporter cells via lentivirus-mediated transduction and observed that overexpression of each Fzd and DVL protein mimicked the effect of Wnt5a by decreasing the WRP reporter signal significantly, whereas overexpression of the Myc eptitope tag as a negative control did not decrease WRP reporter fluorescence (Figure 3E and 3F). These findings suggest that Fzd and DVL family proteins function downstream of Wnt5a to regulate Pdzrn3 degradation.

In addition to Fzd and DVL proteins, several kinases are known to be involved in both canonical and non-canonical Wnt signaling; specifically, GSK3 and CK1 have been reported to phosphorylate Ror receptors (Yamamoto et al., 2007; Grumolato et al., 2010), and Dvl2 and Dvl3 (Bryja, Schulte, & Arenas, 2007; Bryja, Schulte, Rawal, et al., 2007), respectively. Whether these phosphorylation events are required for Wnt5a-dependent regulation of Pdzrn3, however, remains unknown. To address this question, we treated WRP reporter cells with small-molecule inhibitors targeting CK1 (D4476) or GSK3 (CHIR99021). We observed that both treatments significantly reduced Wnt5a-induced GFP-Pdzrn3 degradation (Figure 3G), thus demonstrating a functional role of both CK1 and GSK3 in Wnt5a-Ror-Pdzrn3 signal transduction.

We previously reported that the atypical kinesin Kif26b is another downstream regulatory target of Wnt5a-Ror signaling (Karuna, Susman, & Ho, 2018; Susman et al., 2017). Because Pdzrn3 and Kif26b are both regulated by the Wnt5a-Ror-Dvl axis, we sought to define the epistatic relationship between Pdzrn3 and Kif26b (i.e., whether these two proteins functions in a linear cascade or in parallel branches). To distinguish between these possibilities, we used CRISPR/Cas9 gene editing to generate cells lacking Kif26b and its homolog Kif26a (Kif26a/b dKO cells; sequences in Supplemental Figure 3A), which we previously showed is also a target of Wnt5a-Ror signaling (Karuna et al., 2018), and tested whether the GFP-Pdzrn3 reporter is still degraded upon rWnt5a stimulation. We observed that genetic deletion of *Kif26a* and *Kif26b* does not hinder the ability of rWnt5a to induce GFP-Pdzrn3 degradation; however, there is a slight but significant increase in GFP-Pdzrn3 degradation in the Kif26a/b dKO cells that can be reversed upon re-expression of Kif26b (Figure 3H). In the converse experiment, we again used CRISPR/Cas9 to generate cells lacking *Pdzrn3* and its homolog *Lnx4* (Pdzrn3/4 dKO cells; sequences in Supplemental Figure 3B and 3C), which is structurally very similar to Pdzrn3 (see Figure 6A). We observed that rWnt5a-induced GFP-Kif26b degradation can still occur (Figure 3I) to a large extent. However, deletion of *Pdzrn3* and *Lnx4* did have a slight but significant effect on reducing GFP-Kif26b degradation, which can be alleviated upon re-expression of Pdzrn3. Taken together, these data indicate that Wnt5a regulation of Pdzrn3 does not require Kif26b and vice versa, suggesting that these two targets are epistatically parallel to each other. However, there may be some degree of cross-talk through a currently unknown mechanism.

**Figure 6.**
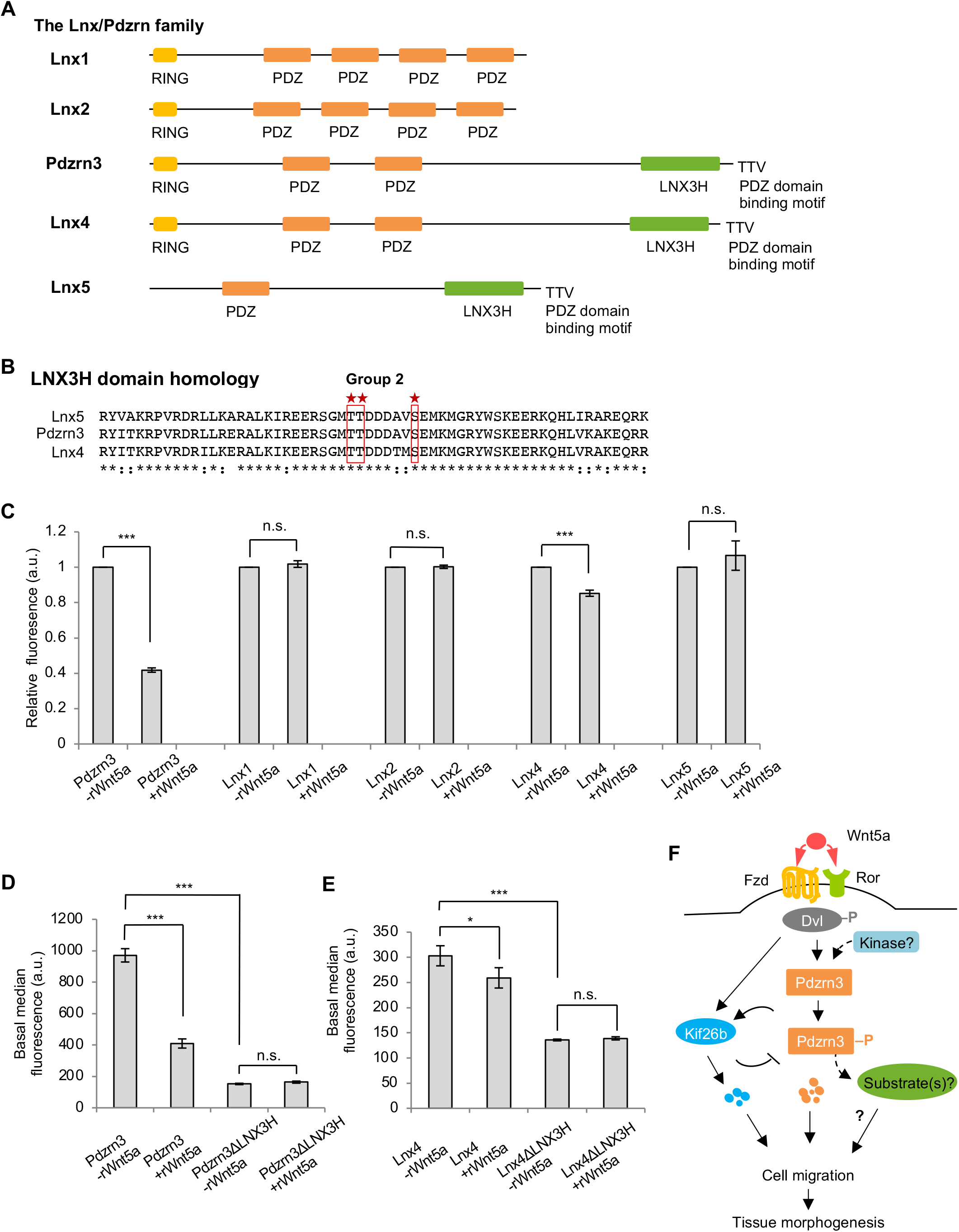
The C-terminal LNX3H domain acts as a general Wnt5a-responsive domain for Pdzrn3 and its homologs. **A)** Schematic of Lnx family members and their conserved domains. Pdzrn3 is structurally most homologous to Lnx4. **B)** Alignment of a portion of the LNX3H domain shared by Pdzrn3, Lnx4, and Lnx5. The Pdzrn3 Group 2 phosphorylation sites identified through our MS screen are conserved (red stars and boxes). **C)** Quantification of the effects of Wnt5a on the steady-state abundance of GFP-Lnx family member reporter cell lines. For clarity and ease of comparison across family members, the median reporter signal for the +Wnt5a condition was normalized to the –Wnt5a condition within the individual Lnx reporter. Error bars represent ± SEM calculated from two (Lnx1, Lnx2, and Lnx5), four (Pdzrn3), or six (Lnx4) cell lines and three technical replicates per line. t-test (unpaired) was performed to determine statistical significance for the following comparisons: +Wnt5a vs. –Wnt5a for each Lnx family member. **D)** Quantification of the effect of LNX3H truncation mutation on Pdzrn3 steady-state abundance. Error bars represent ± SEM calculated from two (Pdzrn3ΔLNX3H) or four (Pdzrn3) cell lines and three technical replicates per line. t-test (unpaired) was performed to determine statistical significance for the following comparisons: WT Pdzrn3 –rWnt5a vs. WT Pdzrn3 + rWnt5a; Pdzrn3 ΔLNX3H –rWnt5a vs. WT Pdzrn3 –rWnt5a; Pdzrn3ΔLNX3H –rWnt5a vs. Pdzrn3ΔLNX3H +rWnt5a. **E)** Quantification of the effect of LNX3H truncation mutation on Lnx4 steady-state abundance. Error bars represent ± SEM calculated from two (Lnx4ΔLNX3H) or six (Lnx4) cell lines and three technical replicates per line. t-test (unpaired) was performed to determine statistical significance for the following comparisons: WT Lnx4 –rWnt5a vs. WT Lnx4 + rWnt5a; Lnx4ΔLNX3H –rWnt5a vs. WT Lnx4 – rWnt5a; Lnx4ΔLNX3H –rWnt5a vs. Lnx4ΔLNX3H +rWnt5a. **F)** Model of Wnt5a-Ror-Dvl-Pdzrn3 signaling. P-values: * = p<0.05, ** = p<0.01, *** = p<0.001.

### Pdzrn3 phosphorylation is required for its Wnt5a-mediated degradation

We next sought to define the structural elements within Pdzrn3 required for its degradation and explore the possible role of phosphorylation in this regulation. Pdzrn3 is a cytosolic protein that contains an N-terminal RING domain that confers its putative E3 ligase activity, two internal PDZ domains that mediate protein-protein interactions, a C-terminal LNX3 homology (LNX3H) domain with no known function, and a C-terminus PDZ domain binding motif (Figure 4A) (Flynn, Saha, & Young, 2011; Sewduth et al., 2014). The six phosphorylation sites identified in our phosphoproteomic analysis cluster into two groups: Group 1 phosphorylation sites (S775, S843, and S845) reside within the linker region between the second PDZ domain and the LNX3H domain, and Group 2 phosphorylation sites (T955, T956, and S962) are located within the LNX3H domain itself. Interestingly, while phosphorylation of Group 1 sites showed only a slight increase at 1 hour and then decreased after 6 hours (compare Figure 1F with dot dashed, dotted, and solid lines in 1G), phosphorylation of Group 2 sites increased significantly after 1 hour of rWnt5a stimulation, prior to Pdzrn3 degradation, and then decreased after 6 hours of rWnt5a stimulation (compare Figure 1F with dashed line in 1G), raising the hypothesis that phosphorylation of these sites, particularly those in Group 2, may be required for Wnt5a-regulation of Pdzrn3 degradation.

To test this hypothesis, we systematically generated phosphoinhibitory mutants (T or S to A substitutions) of all three sites in either Group 1 or Group 2 and examined the effects of these mutations on rWnt5a-induced Pdzrn3 degradation. We observed that while mutation of Group 1 sites had no effect on rWnt5a-induced Pdzrn3 degradation, mutation of Group 2 sites strongly abolished GFP-Pdzrn3 degradation (Figure 4B). To further dissect which specific sites within Group 2 are required for Pdzrn3 degradation, we individually mutated each of the three sites and observed that any of the three single mutations significantly reduced rWnt5a-induced GFP-Pdzrn3 degradation (Figure 4C). To further test whether phosphorylation of these three residues is sufficient to mimic the effect of Wnt5a-Ror signaling on Pdzrn3 degradation, we generated a triple phosphomimetic mutant (T or S to E) and observed that these mutations, in the absence of exogenous rWnt5a stimulation, constitutively decreased the Pdzrn3 reporter signal to a level comparable to that of wild-type Pdzrn3 upon rWnt5a stimulation, and no further degradation was induced by rWnt5 stimulation (Figure 4D). These experiments establish that Wnt5a-dependent phosphorylation of the three Group 2 sites in the LNX3H domain is both required and sufficient to drive Pdzrn3 degradation.

### Wnt5a-directed cell migration requires Pdzrn3 phosphorylation and degradation

We next investigated what the cell biological consequences of Pdzrn3 phosphorylation and degradation might be. Our previous work demonstrated that Wnt5a-Ror signaling can modulate cell migration through regulation of Kif26b abundance (Susman et al., 2017). We wondered if Wnt5a-Ror regulation of effector abundance might be a general paradigm through which Pdzrn3 is similarly controlled. This possibility seemed particularly salient given that others have demonstrated that Pdzrn3 can function as a promigratory factor in cell morphogenetic events, including HUVEC cell migration *in vitro* and neuronal cell positioning *in vivo* (Baizabal et al., 2018; Sewduth et al., 2014). Thus, we hypothesized that Pdzrn3 abundance, directly regulated by Wnt5a-induced phosphorylation, might ultimately serve to regulate cell migration.

To evaluate our hypothesis, we used real-time single cell tracking to first assess the role of the Pdzrn3 protein itself on cell migration. We took advantage of the Pdzrn3 and Lnx4 double knockout cells (Pdzrn3/4 dKO cells), which provided a platform in which we could directly compare the function and regulation of wild-type Pdzrn3 (Pdzrn3 WT cells) and Group 2 phosphoinhibitory site mutant Pdzrn3 (Pdzrn3 phosphoinhibitory cells) through expression of these proteins without potential influence from the structural homolog Lnx4. First, we observed that cells expressing WT Pdzrn3 cells migrated significantly greater distances than Pdzrn3/4 dKO cells (Figure 5A, quantified in Figure 5B), thereby confirming that Pdzrn3 functions as a promigratory factor (Baizabal et al., 2018; Sewduth et al., 2014). Interestingly, we noticed that Pdzrn3 phosphoinhibitory cells migrated not only significantly further than Pdzrn3/4 dKO cells but also significantly further than Pdzrn3 WT cells as well, suggesting that inhibiting Pdzrn3 phosphorylation could potentially enhance cell migration.

We next assayed for the influence of the Wnt5a-Ror-Pdzrn3 axis on cell migration. We observed that while rWnt5a stimulation had no effect on the distance traveled by Pdzrn3/4 dKO cells, it strongly reduced the distance travelled by WT Pdzrn3 cells (Figure 5C, quantified in Figure 5D). Importantly, this Wnt5a effect on cell migration was completely abolished in Pdzrn3 phosphoinhibitory cells. Taken together, these results establish that phosphorylation-dependent degradation of Pdzrn3 is required for Wnt5a to exert its effect on modulation of cell migration.

### The C-terminal LNX3H domain functions as a Wnt5a-responsive domain to regulate protein abundance of Pdzrn3 and related homologs

Pdzrn3 belongs to the Ligand of Numb-X or Lnx family of E3 ligases (Figure 6A). Like Pdzrn3, each Lnx family member possesses an N-terminal RING domain (with the exception of Lnx5) and one to four internal PDZ binding domains. Like Pdzrn3, Lnx4 and Lnx5 each additionally possess a C-terminal LNX3H domain and a C-terminus PDZ domain binding motif (Flynn et al., 2011; Sewduth et al., 2014). Notably, the LNX3H domains of Lnx4 and Lnx5 possess homologous Group 2 phosphorylation sites found in Pdzrn3 (Figure 6B). Based on our finding that these sites regulate Wnt5a-induced Pdzrn3 degradation, we hypothesized that Lnx4 and possibly Lnx5 may also be regulated by Wnt5a signals and that the LNX3H domain may generally function as a Wnt5a-responsive domain. To test this hypothesis, we generated reporter cell lines stably expressing GFP-Lnx1, -Lnx2, -Lnx4, or -Lnx5 fusion proteins and assessed their ability to undergo degradation in response to rWnt5a stimulation. As predicted, when stimulated with rWnt5a, GFP-Lnx1 and GFP-Lnx2, which lack an LNX3H domain, did not degrade, whereas GFP-Lnx4, which has an LNX3H domain, exhibited a modest but significant degradation response (Figure 6C). Interestingly, GFP-Lnx5, which also has an LNX3H domain but lacks a RING domain, was not degraded after rWnt5a stimulation (Figure 6C). We therefore conclude that, like Pdzrn3, Lnx4 is also a target of Wnt5a signaling. Moreover, the Wnt5a responsiveness of Lnx family members correlates with the presence of both LNX3H and RING domains, as the primary difference between Lnx5 and Pdzrn3/Lnx4 is the N-terminal RING domain.

To further test the idea that the LNX3H domain might act as a Wnt5a-responsive domain, we generated truncation mutants of GFP-Pdzrn3 and GFP-Lnx4 lacking this domain. We observed that rWnt5a-induced degradation was completely abolished in these mutant cells (Figure 6D and 6E). In addition, the steady-state fluorescence of unstimulated reporter cells was also substantially reduced (Figure 6D and 6E). These observations suggest that the LNX3H domain of Pdzrn3 and Lnx4 acts not only as a Wnt5a-responsive domain but may do so by regulating overall protein stability, possibly by inhibiting the N-terminal RING domain to prevent auto-ubiqutination and degradation. While the precise mechanism by which the LNX3H domain responds to Wnt5a signals remains unknown and is beyond the scope of this study, our finding defines the LNX3H domain as a bona fide Wnt5a-responsive domain that regulates Pdzrn3 and Lnx4 stability.

## Discussion

In this study, we conducted a whole proteome-scale mass spectrometry screen in primary *Wnt5a* knockout MEFs to identify early and late downstream events driven by Wnt5a-Ror signaling and identified the E3 ubiquitin ligase Pdzrn3 as a regulatory target. Activation of Wnt5a-Ror signaling results in the regulation of Pdzrn3 abundance in a β-catenin-independent manner mediated by a signaling cascade involving Fzd receptors, Dvl scaffolding proteins, GSK3, and CK1 that culminates in UPS-dependent degradation of Pdzrn3. We find that Kif26b is not required for Wnt5a-mediated Pdzrn3 degradation nor is Pdzrn3 required for Kif26b degradation, although there is some potential cross-talk between the two effectors (Figure 6F). Importantly, we determined that the Wnt5a-Ror-Pdzrn3 signaling axis serves to modulate cell migration. Wnt5a-induced Pdzrn3 phosphorylation at three residues on its C-terminal LNXH3 domain is required for its subsequent degradation, which is also required for Wnt5a-Ror signaling to reduce cell migration in NIH/3T3 cells. Thus, the biochemical changes observed in our Wnt5a-Ror signaling cascade connect to a distinct cell biological behavior. Finally, we note that truncation of the LNX3H domain results in constitutive destabilization of Pdzrn3 even in the absence of Wnt5a, suggesting that the LNX3H domain may function as both a Wnt5a-responsive domain and an intrinsic regulator of Pdzrn3 stability. Based on these findings, we propose that the LNX3H domain of Pdzrn3 may function to prevent Pdzrn3 auto-ubiquitination and self-degradation mediated by its RING domain. Prior to Wnt5a stimulation, Pdzrn3 may adopt a “closed” conformation as its C-terminal PDZ domain binding motif interacts with one of its internal PDZ domains to block its E3 ligase activity. Upon Wnt5a stimulation, Pdzrn3 is C-terminally phosphorylated on its LNX3H domain by a yet unidentified kinase to switch the “closed” conformation into an “open” conformation, allowing Pdzrn3 to catalyze the ubiquitination of relevant substrates as well as itself. Notably, this “opened/closed” conformation paradigm has been previously described in other components of Wnt signaling, including Axin and Dvl (Kim et al., 2013; H. J. Lee, Shi, & Zheng, 2015; Qi et al., 2017). Conceivably, the equilibrium between the “closed” and “open” Pdzrn3 could be modulated through either intramolecular interactions within a single Pdzrn3 molecule or through intermolecular interactions between Pdzrn3 dimers or multimers. Future detailed biochemical experiments are required to directly dissect these possibilities as well as evaluate whether Pdzrn3 is phosphorylated by a kinase known to be involved in non-canonical Wnt signaling (such as CK1 or GSK3) or another one, as well as how the kinase itself is regulated by Wnt5a-Ror signaling.

It is well established that several core components of canonical Wnt signaling (e.g., β-catenin, adenomatous polyposis coli (APC), and Axin) are regulated by proteasomal degradation (Choi, Park, Costantini, Jho, & Joo, 2004; Huang et al., 2009; Papkoff, Rubinfeld, Schryver, & Polakis, 1996). The present work, together with other recent studies, establishes that multiple effectors of non-canonical Wnt pathways, including Pdzrn3, Kif26a, Kif26b and Syndecan4, are also subject to regulation by the ubiquitin-proteasome pathway (Carvallo et al., 2010; Karuna et al., 2018; Susman et al., 2017). Collectively, these findings suggest that regulated proteolysis to tune the abundance of downstream effectors and thus, signaling outcomes, may be a conserved paradigm common to both canonical and non-canonical Wnt signaling pathways. This concept will continue to evolve as additional Wnt signaling components are discovered and characterized.

While our study dissects the biochemical regulation of Pdzrn3 by Wnt5a-Ror signaling, previous work by others supports the physiological importance of Pdzrn3 in non-canonical Wnt signaling. One particularly notable study focuses on the role of Pdzrn3 in vascular morphogenesis during embryonic development (Sewduth et al., 2014). In this study, Sewduth et al. identified a binding interaction between Pdzrn3 and Dvl3 via a yeast 2-hybrid screen and a subsequent co-immunoprecipitation, going on to demonstrate that loss of Pdzrn3 *in vivo* results in increased vasculature disorganization in both the embryonic yolk sac and the developing mouse brain. Furthermore, deletion of Pdzrn3 led to decreased persistent directional migration in HUVECs *in vitro*. Importantly, our findings further build upon this model by demonstrating that Wnt5a-Ror signaling can modulate cell migration through Pdzrn3 by triggering its phosphorylation and subsequent degradation. Our study, taken together with existing Pdzrn3 literature, indicates that changes in Pdzrn3 abundance results in non-canonical Wnt signaling defects that can be observed at the molecular, cell, and organismal levels, and supports a physiologically relevant role for Pdzrn3 in Wnt5a-dependent morphogenetic regulation.

The similar means by which Pdzrn3 and Kif26b are regulated indicate that the Wnt5a-Ror pathway has evolved multiple effectors to exert appropriate biological outcomes. Pdzrn3 and Kif26b are regulated by highly similar signaling cascades that uiltize known Wnt signaling components, including Ror receptors, Dvl scaffolding proteins, and GSK3, culminating in UPS-dependent degradation of both effectors. Further, both Pdzrn3 and Kif26b perform related functions at the cell behavioral level. In this study, we describe the mechanism by which Wnt5a-Ror signaling utilizes Pdzrn3 phosphorylation and degradation to modulate NIH/3T3 cell migration. This paradigm is remarkably similar to the one we reported previously, wherein Wnt5a-mediated Kif26b degradation also results in decreases in cell migration as assayed via wound closure in scratch assays (Susman et al., 2017). Our genetic epistasis experiments indicate that Pdzrn3 and Kif26b reside neither upstream nor downstream of each other but do influence each other’s Wnt5a-driven degradation, further suggesting that these two components work in parallel to properly execute signaling funcitons. Thus, Wnt5a-Ror signaling appears to have evolved multiple effectors to ensure tightly coordinated cell biological outcomes. Individual roles for Pdzrn3 and Kif26b, including potential substrates and co-effectors, should be examined in future studies.

The lack of quantitative and reliable readouts for Wnt5a-Ror signaling has been a major limitation in the field. We leveraged our discovery of Pdzrn3 and its regulation by Wnt5a-Ror signaling to develop a new flow cytometry-based reporter that enables sensitive and quantitative detection of pathway activity in live cells. In addition to dissecting the mechanisms that mediate Pdzrn3 degradation, this reporter assay could also be utilized to interrogate other biochemical steps in the pathway upstream of Pdzrn3, understand various disease-associated mutations, and serve as an important platform for high throughput screening of small molecules that target Wnt5a-Ror-driven developmental disorders and cancers.

## Materials and methods

### Cell lines

Primary MEFs were isolated directly from mouse embryos as described (Ho et al., 2012) and used within 3 passages. NIH/3T3 Flp-In (R76107, Thermo Fisher Scientific) cells were purchased and were not re-authenticated; cells tested negative for mycoplasma contamination using the Universal Mycoplasma Detection Kit (30-1012K, ATCC). All cell lines were cultured at 37C and 5% CO2 in Dulbecco’s Modified Eagles Medium (MT15017CV, Corning) supplemented with 1x glutamine (25-005-CI, Corning), 1x penicillin-streptomycin (30-002-CI, Corning) and 10% fetal bovine serum (16000069, Thermo Fisher Scientific).

### TMT/MS3 proteomic screen

Primary *Wnt5a*^-/-^ MEFs (derived and pooled from three different E12.5 *Wnt5a*^-/-^ embryos) were seeded in six 10-cm plates at 50% confluency 3 days before rWnt5a stimulation (day 0), such that cells would be fully confluent for 2 days. On the day of stimulation (day 3), cells in each 10-cm plate were treated either with rWnt5a (100ng/mL final concentration) for 1h or 6hr, or with the control buffer (1x PBS, 0.1% bovine serum albumin, 0.5% w/v CHAPS) for 6hr. The entire stimulation experiment was conducted in two independent replicates. At the end of the Wnt5a stimulation time course, cells were washed once with ice-cold PBS and plates were scraped into 1 mL of ice-cold lysis buffer (8 M urea, 75 mM NaCl, 50 mM Tris pH 8.2, 1 mM NaF, 1 mM β-glycerophosphate, 1 mM Na_3_VO_4_, 10 mM Na_4_P_2_O_7_, 1 mM PMSF, and Complete protease inhibitor (-EDTA, Roche)). Cells were homogenized by pipetting up and down using a P-1000 and then sonicated in a Bioruptor (17 x 30s ON/OFF cycles). Cell lysates were then centrifuged at 40,000 RPM for 20 min at 4C. The clarified high-speed supernatants were collected, snap frozen in liquid nitrogen and stored at −80C until the TMT/MS3 analysis was performed. Protein concentrations were determined using BCA reagents (Pierce) and normalized.

To perform the TMT/MS3 screen, tryptic peptides were prepared from whole cell lysates and the peptide mixtures from the different experimental conditions were labeled with the six TMT reagents, such that reporter ions at m/z of 126, 127, 128, 129, 130 and 131 would be generated in the tandem spectrometry. Phosphopeptides were enriched by TiO_2_ chromatography. Liquid chromatography, MS3 tandem mass spectrometry and data analysis were carried out as previously described (McAlister et al., 2014; Paulo et al., 2015; Ting et al., 2011).

### Cloning of mouse *Pdzrn3, Lnx1, Lnx2, Lnx4*, and *Lnx5* cDNA

For cloning of mouse Pdzrn3 cDNA, a first strand cDNA pool was generated from MEF total RNA Maxima H Minus reverse transcriptase and oligo dT primers according to manufacturer’s instructions (EP0751, ThermoFisher Scientific). This cDNA library was then used as template for PCR amplification of the *Pdzrn3* open reading frame with the following primers, forward: gatcGGCCGGCCtACCatgggtttcgagttggatcgc; reverse: gatcGGCGCGCCTTATACAGTAGTCACCGACAGGAA. The PCR product was subcloned into a modified pCS2+ vector using the FseI and AscI restriction sites. The entire *Pdzrn3* open reading frame was confirmed by Sanger sequencing.

For cloning the Lnx1, Lnx2, Lnx4, and Lnx5 cDNAs, the same workflow was used, except that E14.5 mouse brain RNA was used to generate the first strand cDNA pool. The following primers were used to PCR amplify and subclone the respective cDNAs: mLnx1 forward, gatcGGCCGGccTACCatgaaccaaccggaccttgcagat; mLnx1 reverse, gatcGGCGCGCCTTATAAAAAAGTACCAGGCCAAGAAG; mLnx2 forward, gatcGGCCGGccTACCatgggaacaaccagtgacgagatgg; mLnx2 reverse, gatcGGCGCGCCCTATACGAGGCTGCCTGGCCAGCAG; mLnx4 forward, gatcggccggccTaccATGGGCTTCGCTTTGGAGCGTCTC; mLnx4 reverse, gatcGGCGCGCCtcaTACGGTGGTCACCGACAGAAAGGC; mLnx5 forward, gatcGGCCGgCCTACCatgggatgtaatatgtgtgtggtc; mLnx5 reverse, gatcGGCGCGCCTCAGACAGTGGTGACAGAGAGCAG. All constructs were confirmed by Sanger sequencing.

### Antibodies

Antibodies against Ror1, Ror2, and Kif26b were described previously (Ho et al., 2012; Susman et al., 2017). The following antibodies were purchased: rabbit anti-Dvl2 (#3216, Cell Signaling) and mouse anti-α-tubulin (clone DM1A, #ab7291, Abcam).

Initial analyses of Pdzrn3 were conducted using a commercial antibody (SC-99507, Santa Cruz Biotechnology); however, the antibody was discontinued and all subsequent analyses (including all data presented in this paper) were conducted using anti-Pdzrn3 antibodies produced in-house. To generate anti-Pdzrn3 antisera, rabbits were immunizing with a mixture of two different antigens: 1) a synthetic peptide with the sequence LLTHGTKSPDGTRVYNSFLSVTC, conjugated to keyhole limpet hemocyanin (77600, ThermoFisher Scientific), and 2) a maltose binding protein N-terminally fused to a Pdzrn3 protein fragment extending from amino acids 902 to 1063, recombinantly expressed in and purified from E. coli. Antibodies were affinity purified from antisera over a column with a full-length recombinant Pdzrn3 protein covalently immobilized to Sepharose beads (AminoLink Plus, 20501, ThermoFisher Scientific). Full length Pdzrn3 was expressed in insect cells using the Bac-to-Bac baculovirus expression system (10359016, ThermoFisher Scientific); the protein was insoluble and was purified under denatured conditions using 5.5M guanidinium hydrochloride, coupled to AminoLink Plus Resin, and renatured by gradually removing guanidinium hydrochloride.

### Western blotting

Protein lysates for SDS-PAGE and western blotting were prepared in 1x - 2x Laemmli sample buffer or LDS sample buffer (Life Technologies). Protein lysates used for Kif26b western blotting were not heated, as the Kif26b signal weakens substantially after heating, likely due to heat-induced protein aggregation (Susman et al., 2017). All other protein lysates were heated at 90C for 5 min before SDS-PAGE and western blotting.

Quantitative western blotting was performed using the Odyssey infrared imaging system (Li-Cor Biosciences) according to the manufacturer’s instructions. The median background method was used with a border width of two pixels on all sides around the perimeter of the area being quantified. Non-saturated protein bands were quantified by using Odyssey software with the gamma level set at 1.

### Generation of stable NIH/3T3 cell lines

To construct the GFP-Pdzrn3 expression plasmid, the eGFP open reading frame was first subcloned into pENTR-2B (Life Technologies), and the full-length mouse Pdzrn3 open reading frame was subcloned in frame to the C-terminus of GFP. The resulting construct was verified by sequencing and then recombined with the pEF5-FRT-V5 vector (Life Technologies) using LR Clonase (Life Technologies) to create pEF5-GFP-Pdzrn3-FRT. The pEF5-GFP-Pdzrn3-FRT plasmid was used to generate stable isogenic cell lines using the Flp-In system and Flp-In NIH/3T3 cell line (Life Technologies). DNA transfection was performed in 10-cm plates with GenJet In Vitro Transfection Reagent (SL100488; SignaGen Laboratories). Cells that stably integrate the Flp-In constructs were selected using 200μg/ml hygromycin B and expanded. Cell lines expressing phosphoinhibitory or phosphomimetic Pdzrn3, Lnx1, Lnx2, Lnx4, Lnx5, Pdzrn3ΔLNX3H, and Lnx4ΔLNX3H were similarly created by cloning the open reading frame to the C-terminus of GFP in frame and conducting the workflow described above.

### Lentivirus-mediated protein overexpression

Recombinant lentiviruses were generated using the pLEX_307 (for all Fzd and DVL constructs) vectors, which uses the EF1 promoter to drive transgene expression. pLEX_307 was a gift from David Root (Addgene plasmid # 41392). The human DVL1 and DVL3 open reading frames were cloned by PCR from a HeLa cell cDNA pool using the following primers; hDVL1 forward, gatcGAATTCCACCatgggcgagaccaagattatctac; hDVL1 reverse, gatcGGCGCGCCTCACATGATGTCCACGAAGAACTC; hDVL3 forward, TTCAGGCCGGCCTACCATGGGCGAGACCAAGATCATCTAC; hDVL3 reverse, GAGGCGCGCCTCACATCACATCCACAAAGAACTC. Similarly, the human DVL2 open reading frame was cloned by PCR from a separate HeLa cDNA pool. The following primers were used: hDvl2 forward, gcggcggcgGcCgGccaatggcgggtagcagcactggggg; hDVL2 reverse, gtcgacgGgCGcgcctacataacatccacaaagaactcg. The mouse Fzd1 and Fzd7 open reading frames were PCR amplified from Addgene plasmids #42253 and 42259 (gifts from Jeremy Nathans), respectively, using the following primers: mFzd1 forward, gatcggccggcctaccatggctgaggaggcggcgcctag; mFzd1 reverse, gatcggcgcgccTCAGACGGTAGTCTCCCCCTGTTTG; mFzd7 forward, gatcggccggcctaccatgcggggccccggcacggcggcg; mFzd7 reverse, gatcggcgcgccTCATACCGCAGTTTCCCCCTTGC. The mFzd2 open reading frame was cloned via PCR from mouse brain via the following primers: mFzd2 forward, gatcggccggcctaccatgcgggcccgcagcgccctg; mFzd2 reverse, gatcggcgcgccTCACACAGTGGTCTCGCCATGC. The open reading frames of all lentiviral constructs were verified by sequencing. Lentiviruses were packaged and produced in HEK293T cells by co-transfection of the lentiviral vectors with the following packaging plasmids: pRSV-REV, pMD-2-G and pMD-Lg1-pRRE (gifts from Thomas Vierbuchen). 0.75ml or 0.25 ml of the viral supernatants was used to infect GFP-Pdzrn3 reporter cells seeded at 20% confluency in 24-well plates. Puromycin selection (0.002 mg/ml) was carried out for three days. Cells from the viral titer that killed a large proportion of cells (60-90%) were expanded and used for flow cytometry; this ensured that the multiplicity of infection (MOI) is ∼1 for all cell lines used in the experiments. This same workflow was utilized to establish GFP-Pdzrn3 and GFP-Kif26b reporters in Kif26a/b dKO cells and Pdzrn3/4 dKO cells, respectively; in lieu of puromycin selection, GFP-positive cells were sorted (MoFlo Astrios Cell Sorter, Beckman Coulter, 488nm laser) and expanded prior to degradation analysis.

### Generation of double knockout cell lines

*Kif26b* knockout cells were previously described (the mutant clone with +1 and −13 frameshifts, generated using sgRNA 1; (Susman et al., 2017)). This *Kif26b* mutant clone was subject to a second round of mutagenesis to knock out *Kif26a* via CRISPR/Cas9-mediated genome editing according to (Ran et al., 2013). Briefly, a modified version of LentiCRISPR V2 (Addgene #52961), in which the puromycin selection cassette was modified with a blasticidin selection cassette, was used to generate lentiviruses expressing small guide RNAs (sgRNAs) with the following sequence: GCTCGTGGAGCTAAAACGAC. In wild-type NIH/3T3 cells, Pdzrn3 was similarly targeted using the following sequence: AGCTGCCCCGCGCGTTGTCG. Following lentivirus infection, cells were passaged for 5 days to allow time for mutagenesis to occur. Cells were subsequently selected using blasticidin (0.002mg/mL) in the case of *Kif26a* mutagenesis, or puromycin (0.002mg/mL) in the case of *Pdzrn3* mutagenesis. Individual cell clones were picked from cell populations targeted with each of these sgRNAs, expanded and then validated by deep sequencing the relevant genomic regions amplified by PCR.

To generate *Pdzrn3/Lnx4* double knockout cells, *Pdzrn3* knockout NIH/3T3 cells were electroporated with CRISPR/Cas9 ribonucleoprotein complexes targeting *Lnx4* using the following gRNA sequence: GCCAACAUCGGCAUGACUCGUUUUAGAGCUAUGCU. 24 hours after electroporation, cells were subjected to fluorescence activated cell sorting (MoFlo Astrios Cell Sorter, Beckman Coulter, 561nm laser) to plate individual cells in 96-well plates; cells were allowed to recover for two weeks prior to expansion and validation of mutations via deep sequencing the relevant genomic regions amplified by PCR.

### Recombinant proteins and inhibitors

The following recombinant proteins and drugs were purchased: human/mouse Wnt5a (654-WN-010, R&D Systems); Wnt-C59 (C7641-2s; Cellagen Technology); epoxomicin (A2606, ApexBio); PYR-41 (B1492, ApexBio); MLN4924 (I50201M, R&D systems); mouse Dkk-1 (5897-DK-010, R&D Systems); IWR-1-endo (B2306, ApexBio); D4476 (A3342, ApexBio); and CHIR99021 (A3011, ApexBio).

### Reverse transcription and qPCR

Total RNA was isolated from *Wnt5a* KO MEFs stimulated with rWnt5a for 0, 1, or 6 hours using the RNeasy Plus Mini Kit (Qiagen, #74134), and cDNA was synthesized using QuantiNova Reverse Transcription Kit (Qiagen, #205411), both according to the manufacturer’s instructions. The cDNA was the source of input for qPCR, using QuantiNova SYBR Green PCR Kit (Qiagen, #208054). The following qPCR primer pairs were used: mPdzrn3 forward, CTGCGCTACCAGAAGAAGTTC; mPdzrn3 reverse, TCCATCTTGATTGTCCACACAG; mGapdh forward, AGGTCGGTGTGAACGGATTTG; mGapdh reverse, TGTAGACCATGTAGTTGAGGTCA.

### Flow cytometry

NIH/3T3 cells were plated at a density of 0.09-0.095M/well in 48-well plates either directly in complete media containing Wnt-C59 (10nM) or in complete media and later changed to complete media containing Wnt-C59 24 hours after plating; all rWnt5a stimulations and inhibitor pretreatments and treatments were conducted in the presence of Wnt-C59. 48 hours after plating, cells were stimulated with rWnt5a for 6 hours. For inhibitor treatments, cells were pretreated with the appropriate inhibitor for 1 hour prior to rWnt5a treatment for 6 hours in the presence of the same inhibitor. Cells were then harvested, resuspended in PBS + 0.5% FBS and analyzed using a flow cytometer (Becton Dickinson FACScan, 488nm laser). Raw data were acquired with CellQuest (Becton Dickinson) and processed in FlowJoX (Treestar, Inc). Processing entailed gating out dead cells, calculation of median fluorescence, percent change of medians, and overlay of histograms. Dose-response curves based on percent change were fitted in Prism (GraphPad Software).

### Live cell imaging and 2D cell migration

Pdzrn3/Lnx4 knockout cells, Pdzrn3 WT cells, and Pdzrn3 phosphoinhibitory cells (all NIH/3T3 cells) were cultured in complete media or in Wnt-C59 containing media for 72 hours (for experiments involving rWnt5a stimulation). Cells were subsequently plated at a density of approximately 0.01M cells per 24 well plate for live cell imaging. rWnt5a treatment was initiated immediately prior to imaging. Multipoint time lapse images were collected every 10 minutes for 20 hours on an Andor Dragonfly spinning disc confocal system in a humidity controlled chamber at room temperature (37C). Cell migration was tracked using the ImageJ manual tracking plugin, and cells that divided, moved out of frame, or died were excluded from further analysis. Total distance traversed was calculated using the ImageJ Chemotaxis tool plugin. Statistical analysis was done using Prism 8 (GraphPad Software).

## Acknowledgements

We thank the members of the Ho and Jao labs at UC Davis for their input and discussions. We thank Karl Willert for careful reading of the manuscript. We acknowledge Bridget McLaughlin and Jonathan Van Dyke at the UC Davis Cancer Center Flow Cytometry core for their training and technical assistance (supported by NCI P30 CA093373). We thank Mikaela Louie and Alec Konopelski Snavely for their assistance with MatLab. We also thank Ryan Toedebusch and Christine Toedebusch for discussions about CRISPR targeting strategies as well as Xueer Jiang, Jose Uribe Salazar, and Megan Dennis for their assistance with CRISPR mutation analysis. This work was supported by National Institutes of Health grant 1R35GM119574 and American Cancer Society grant IRG-95-125-13 to H.H. Ho, and National Institutes of Health grant GM67945 to S. P. Gygi.

